# Visuospatial computations vary by category and stream and continue to develop in adolescence

**DOI:** 10.1101/2025.01.14.633067

**Authors:** Jewelia K. Yao, Justin Choo, Dawn Finzi, Kalanit Grill-Spector

**Affiliations:** Department of Psychology, Stanford University, Stanford, CA 94305; Wu Tsai Neuroscience Institute, Stanford University, Stanford, CA 94305; Department of Symbolic Systems, Stanford University, Stanford, CA, 94305

## Abstract

Reading, face recognition, and navigation are supported by visuospatial computations in category-selective regions across ventral, lateral, and dorsal visual streams. However, the nature of visuospatial computations across streams and their development in adolescence remain unknown. Using fMRI and population receptive field (pRF) modeling in adolescents and adults, we estimate pRFs in high-level visual cortex and determine their development. Results reveal that pRF location, size, and visual field coverage vary across category, stream, and hemisphere in both adolescents and adults. While pRF location is mature by adolescence, pRF size and visual field coverage continue to develop – increasing in face-selective and decreasing in place-selective regions – alongside similar development of category selectivity. These findings provide a timeline for differential development of visual functions and suggest that visuospatial computations in high-level visual cortex continue to be optimized to accommodate both category and stream demands through adolescence.

## Introduction

Across development, the ability to perceive faces, words, bodies, and places is crucial for key behaviors like social interactions, reading, and navigation, and depends on computations by neurons throughout visual cortex^1–5^. In humans, perception of these categories is enabled by computations in category-selective regions across three distinct visual processing streams emerging from early visual cortex (EVC; V1-V3): the ventral stream, which extends ventrally from occipital to temporal cortex and is involved in visual recognition^1,5^, the dorsal stream, which runs through superior occipital-parietal cortex and is engaged in spatial navigation and attention^1,5^, and the lateral stream, which extends from lateral occipitotemporal cortex through the superior temporal sulcus (STS) and is involved in dynamic, action, and social perception^2–4,6^. While many studies examined functional differences among the three pathways, few studies have investigated how basic visual properties such as receptive fields (RFs) – the part of visual space processed by a neuron^7^ and population receptive fields (pRFs) – the portion of visual space processed by the population of neurons in a voxel^8^– differ within and across streams. Even less is known about how pRFs in category-selective regions develop, particularly in adolescence when visual behaviors like face recognition and reading, and the visual areas supporting them, are still developing^9–11^. Given these gaps in knowledge, we ask: (1) How do pRFs in high-level category-selective regions differ across the ventral, dorsal, and lateral streams? (2) Do pRFs develop during adolescence?

In each visual area, pRFs are organized systematically, tiling the visual field; this is referred to as visual field coverage (VFC) of an area^12^. While much is known about VFC and pRF properties in early retinotopic areas^13,14^, pRFs in high-level category-selective regions in the ventral, lateral, and dorsal streams have been less studied, partly because traditional pRF mapping experiments used flickering checkerboards designed to activate early and intermediate, rather than high-level visual areas^8,15–20^. Nonetheless, more recent studies in adults have employed pRF mapping stimuli that include shapes, objects, faces, and colors that drive neurons in high-level regions, consequently enabling the estimation of pRFs in category-selective regions^12,21–26^. Findings of differential retinotopic biases and pRFs in high-level visual cortex led researchers to hypothesize about the origin of these differences.

With respect to category, researchers have hypothesized that fixation patterns on different categories have systematic retinotopic biases that are reflected in retinotopic biases of the respective category-selective region. For example, adults fixate on faces and words in order to recognize faces and read, respectively, and pRFs and VFC in adults’ ventral face- and word-selective regions are concentrated around the center of gaze (fovea) ^22,27,28,28,29^. In general, when people fixate on faces, bodies will be below the face, and this is mirrored in the lower-visual field bias of pRFs of ventral body-selective regions^30^. Additionally, in the real world, places encompass the entire visual field, and pRFs and VFC in ventral place-selective areas extend into the periphery^22,28,29,31,32^. The category hypothesis thus suggests that pRF differences across category-selective regions are driven by unique spatial configurations and distinct viewing patterns that are associated with different categories^24,28,29,33,34^.

With respect to streams, researchers have hypothesized that different streams have distinct computational objectives or functions^1–5^ that utilize visual information from different parts of the visual field. Some studies suggest differences in upper/lower visual field biases across streams with ventral stream pRFs processing the upper visual field and dorsal stream pRFs processing the lower visual field^35,36^. Other studies suggest differences in eccentricity biases across streams whereby pRFs and VFC in ventral face selective regions are centrally biased, but those in lateral face-selective regions extend to the periphery^2,22^. Thus, the stream hypothesis predicts that pRFs will systematically vary across streams. Nonetheless, the category and stream hypotheses are not mutually exclusive, as pRFs and VFC may vary by both category and stream.

The category and stream hypotheses offer frameworks for how pRF properties and VFC might differ across visual cortex in adults, but they do not make predictions regarding developmental trajectories. Adolescence, the period between ages 10 to 19 years, presents a unique developmental window as crucial visual behaviors, like face recognition, reading and spatial attention, along with the underlying category selectivity in face- and word-regions, are still developing^10,37–40^. Currently, we have no knowledge of retinotopic development after age 12^27,41,42^.

Thus, understanding how pRFs may develop during this period will provide important insights into the timeline of development of basic visual functions. We consider two possibilities regarding pRF development: One possibility is that as category selectivity develops into adolescence^10,37–39^, pRF properties in high-level visual areas will also continue to develop into adolescence. This hypothesis predicts that during adolescence, pRF properties and VFC in category-selective regions will be different from that of adults, and is supported by studies finding that pRFs in pFus-faces and pOTS-word continue to develop from age 5 to adulthood^27^. Alternatively, as pRFs perform basic spatial computations on visual inputs, they may mature before higher-level, category computations, and thus may be fully developed by adolescence. This hypothesis predicts no significant difference in the properties of pRFs and VFC in category-selective regions between adolescents and adults and is supported by work demonstrating that pRF properties and VFC of early visual areas (V1-V3) are adultlike as early as 5 to 7 years of age^27,41,42^. Of course, it is also possible that differential development occurs whereby pRFs/VFC in some streams or category-selective regions mature by adolescence while others continue to develop.

## Results

### Toonotopy drives high-level category-selective regions in adolescents and adults

To test these hypotheses, 15 adolescents (ages 10 - 17; 9 females, 6 males) and 27 adults (ages 22 - 32; 13 females, 14 males) participated in two fMRI experiments, one to map pRFs using sweeping bars with cartoons (Toonotopy, Finzi 2021, Fig. 1A) and another to identify category-selective regions (functional localizer, Fig. 2A). To map pRFs, participants completed four runs of Toonotopy while fixating and performing a color task on fixation (Fig. 1A; Finzi et al., 2021). We model each voxel’s pRF with a 2-D Gaussian defined by three main parameters – its location (*x,y*) in the visual field, size (*σ*) or the area of the visual field it processes, and a compressive nonlinearity (*n)* (Fig. 1B, Kay 2013). Using pRF estimates, we generated polar angle, eccentricity, and size maps in every participant.

**Figure 1.**
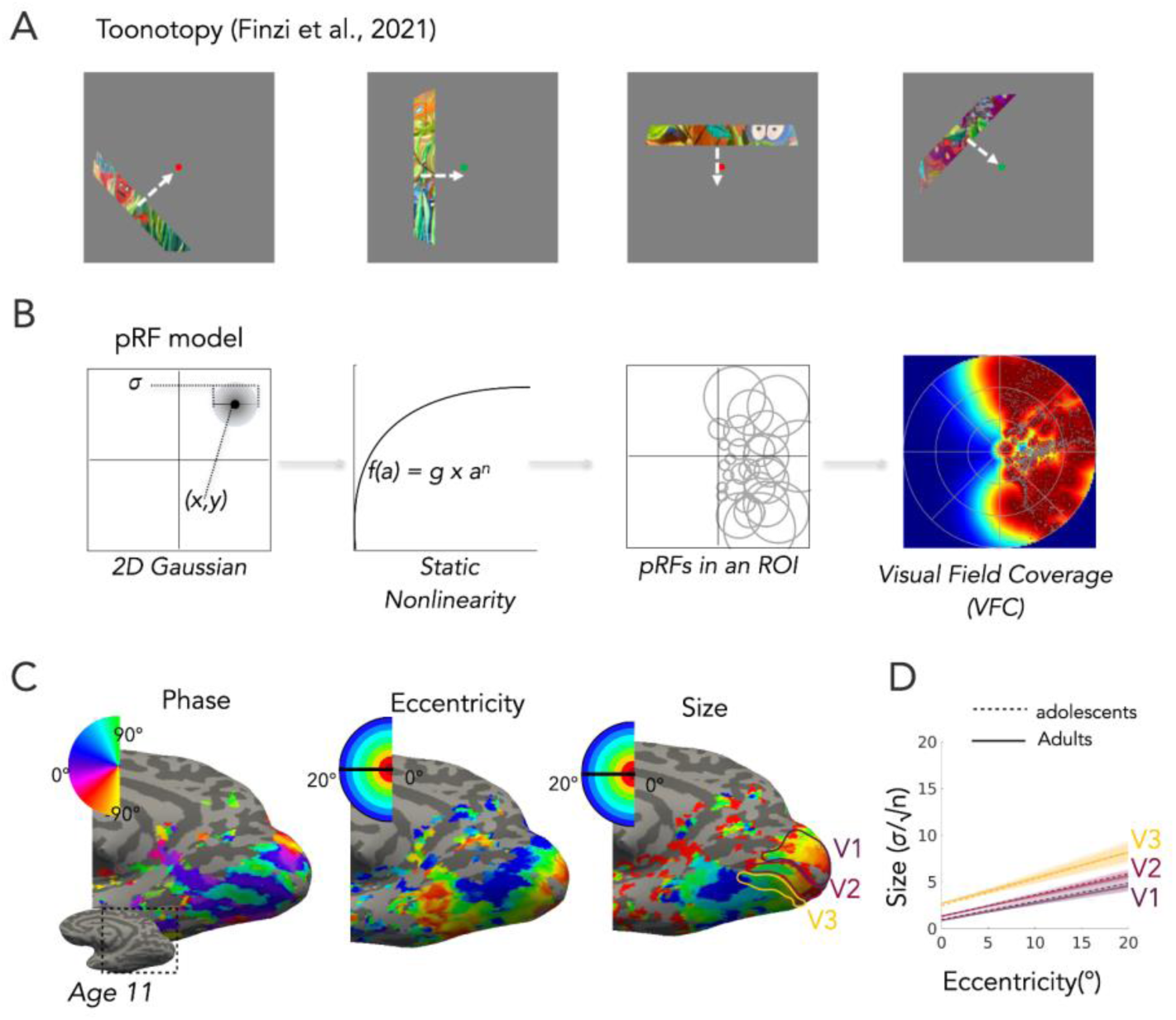
*Toonotopy experiment* (A) Toonotopy stimuli from Finzi et al. (2021) features a bar with colorful cartoon images of faces, words, bodies, places, and objects that change at 8Hz sweeps across a gray background at 4 angles (0°, 45°, 90°, 135°) each in 2 directions. Participants fixated at the center dot and indicated when the dot changed colors. (B) Population receptive field compressive spatial summation (pRF CSS) model^19^. Left two panels show a single pRF with parameters of location (x,y) and size (*σ*) modeled by a 2D Gaussian followed by a compressive nonlinearity, used to model the voxel’s response. Middle right panel shows schematic of the pRF distribution within an ROI, and the rightmost panel depicts the visual field coverage of all pRFs in left V1 ROI in an example 11-year-old. (C) Phase, eccentricity, and size maps in an example adolescent (age 11) with V1 (purple), V2 (magenta), and V3 (gold) borders illustrated on the size map. All participants - Supplementary Fig. 2.(D) pRF size versus eccentricity relationship is similar across adolescents (dotted line) and adults (solid line) in early visual cortex (V1 - V3).

**Figure 2.**
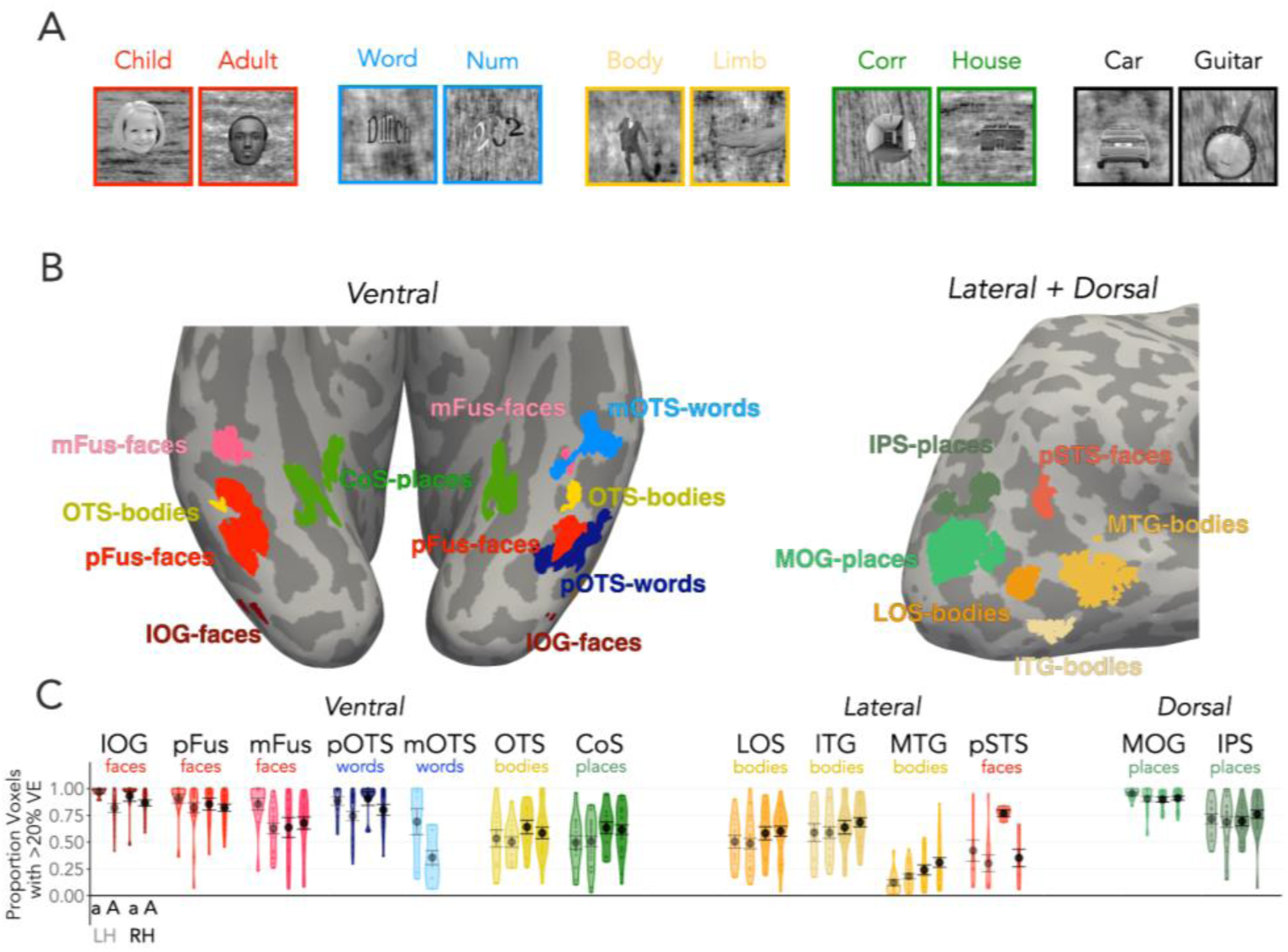
*Category-selective regions are modulated by the Toonotopy experiment* (A) Functional localizer (fLOC): Example stimuli of faces (children, adults; red), characters (words, numbers; blue), bodies and limbs (yellow), places (corridors, houses; green), and objects (car, guitar; black) from the fLOC experiment). (B) We identified in each participant category-selective functional regions of interest (ROIs) from the fLOC experiment, ROIs are labeled by preferred category and anatomical locations. 7 ROI were in the ventral stream (IOG-faces, pFus-faces, mFus-faces, pOTS-words, mOTS-words, OTS-bodies, CoS-places), 4 ROIs in the lateral stream (pSTS-faces, LOS-bodies, ITG-bodies, MTG-bodies), and 2 ROIs in the dorsal stream (MOG-places, IPS-places). *Left:* bilateral ventral ROIs in an example 17-year-old. *Right:* dorsal and lateral ROIs in the right hemisphere in an example 11-year-old. Left hemisphere dorsal and lateral ROIs are the same as the right hemisphere ROIs. (C) Violin plots of proportion of voxels with greater than 20% variance explained in each pRF in category-selective ROIs in the left hemisphere (light) and right hemisphere (dark) ventral, lateral, and dorsal streams in adolescents (a) and adults (A). Black circle: mean. Error bars: ± SE (standard error of the mean). Each dot is a participant.

We first assessed if adolescents have the expected retinotopic maps^8,13,15,31^. In every adolescent, we find the expected polar angle map with mirror reversals of the upper and lower visual field representations beginning in the calcarine sulcus (Fig. 1C - Phase, example participant, Supplementary Fig. 1, all participants). Adolescents also exhibit the expected occipital eccentricity maps showing a gradient of foveal to peripheral representations beginning at the occipital pole and moving anteriorly, as well as a second temporal eccentricity map showing a gradient of foveal to peripheral representations from lateral to medial ventral temporal cortex (VTC, Fig. 1C – Eccentricity). Additionally, adolescents have characteristic pRF size maps with pRFs increasing from small to large beginning at the occipital pole and moving anteriorly along the calcarine as well as from the occipital cortex to ventral temporal cortex (Fig. 1C - Size). As expected, pRF x-position, y-position, eccentricity, and size in V1, V2, and V3 do not develop from adolescence to adulthood (Supplementary Table 1). For all participants, we quantified the well-established relationship between pRF size and eccentricity across early visual areas^8,12,19^. In both adolescents and adults, pRF size increases approximately linearly with eccentricity, and the slopes of this relationship increase from V1 to V3 (Fig. 1D, linear mixed model, LMM, *pRF Size ∼ Eccentricity x ROI (V1/V2/V3) x Age Group x Hemisphere + (1*|*Participant)*), main effect of ROI: p = 7.68*10^-25^, F(2,165) = 79.22). We find no significant difference between age groups in pRF size vs. eccentricity slopes (p = 0.89, F(1,33) = 0.02) or intercepts (p = 0.59, F(1,33) = 0.29) or any interactions with age group (p’s > 0.18, F’s < 1.70; all stats, Supplementary Table 2). Together, these results show that Toonotopy can be used to map pRFs in adolescents.

Using the category localizer experiment (Fig. 2A), we define face, word, body, and place functional regions of interest (ROIs) in each of these participants (Fig. 2B) and estimated pRFs in each ROI). ROIs were found in both hemispheres and in most participants (Supplementary Table 3) except for mOTS-words (mostly left hemisphere), pSTS-faces (mostly adults), and MTG-bodies (mostly adults) As prior research combined the dorsal and lateral stream into a single dorsal stream (e.g. Hasson 2003; Silson 2013; 2016) and because streams differ in the number and type of category-selective regions (Fig. 2B) – e.g., dorsal stream contains only one category (places) – throughout this study we statistically compare the ventral stream and a combined dorsal-lateral stream.

We tested whether Toonotopy drives category-selective regions by evaluating the proportion of voxels in each ROI for which the pRF model explained more than 20% variance during the Toonotopy experiment (Fig. 2C & 2D). We chose this threshold as it is typically used in the field (e.g., Finzi et al., 2021). In the ventral and dorsal streams, a majority (∼80%) of voxels are driven by Toonotopy (Fig. 2C). In the MTG and pSTS MPM ROIs in the lateral stream, fewer voxels (20-60%) are driven by Toonotopy (Fig. 2D). Indeed, the proportion of voxels with greater than 20% variance explained differs significantly across category, stream, and hemisphere (*proportion voxels (R^2^>0.2))∼ Age Group x Stream x Category x Hemisphere +(1|Participant);* category x stream x hemi interaction: p = 0.01, F(2, 681.59) = 4.83; ; all stats, Supplementary Table 4). Additionally, adolescents have significantly more voxels driven by the Toonotopy experiment in face-selective regions, but not other category-selective regions compared to adults (significant age group by category interaction (p = 0.01, F(683.89,2) = 4.64, LMM, Supplementary Table 4; (post-hoc t-test on faces: p = 0.02, t(252) = 3.12). Overall, while there is regional variability in the proportion of voxels with greater than 20% variance explained, many of the voxels in category selective regions are driven by the Toonotopy experiment.

### pRFs vary by stream, category, and hemisphere and develop in size but not location

To assess how pRFs properties may vary by processing stream and category, we examined the distribution of pRF centers and their sizes for each ROI in adolescents (Fig 3A - *Top*) and adults (Fig. 3A - *Bottom*). Across all ROIs and groups, pRF centers lie in the contralateral visual field with no significant differences across age groups (Supplementary Fig. 5; LMM: *pRF x-position ∼ Age Group x Stream x Category x Hemisphere +(1|Participant)*, main effect of hemisphere: p = 2.73*10-^230^, F(1, 688.85) = 2476.21 all stats, Supplementary Table 5). However, pRF distributions appear to differ systematically across both category and stream. For instance, pRFs of ventral face ROIs (IOG/pFus/mFus) are concentrated within the central 5°, those of ventral body ROIs (OTS) span the central 10°, and pRFs in lateral face (pSTS) and body ROIs (LOS, ITG, MTG) extend more peripherally up to 20° (Fig 3A). These qualitative observations align with our hypotheses, which make different predictions about a region’s pRF distributions in the center versus periphery or upper versus lower visual field depending on its processing stream or category selectivity. To qualitatively test these predictions, we use linear mixed models (LMMs) to compare pRF parameters across age group (adolescents/adults), streams (ventral/dorsal-lateral), categories (faces, bodies, places), and hemispheres (right/left). To include word-selective ROIs, which were only found in the ventral stream, we used a second, ventral LMM, comparing age group, category, and hemisphere across ventral stream ROIs.

**Figure 3.**
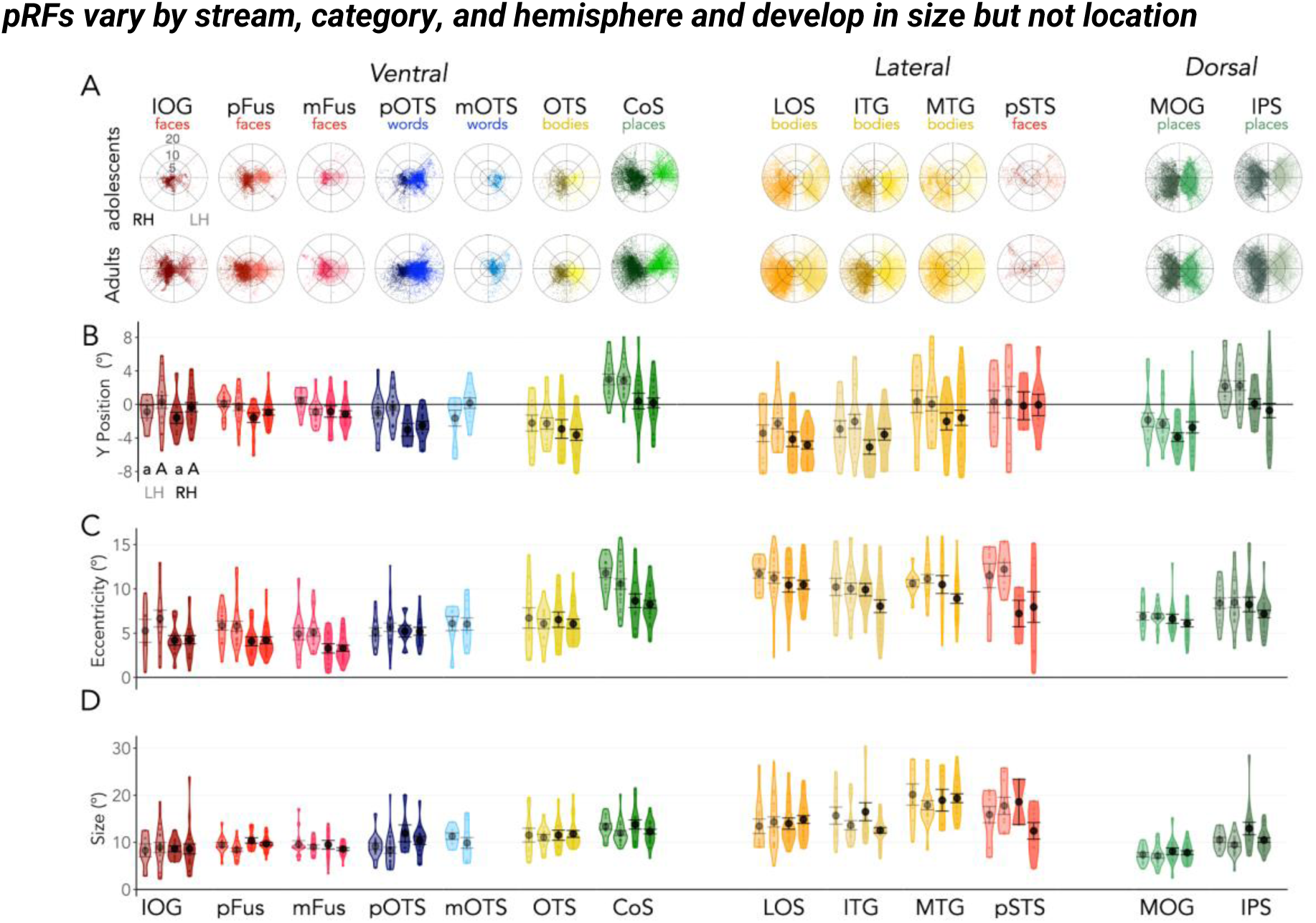
*pRF properties in high-level category selective regions vary across category and stream.* (A) *Top:* pRF center polar plots for right hemisphere (dark) and left hemisphere (light) face-selective (reds; IOG, pFus, mFus, pSTS), word-selective (blues; pOTS, mOTS), bodypart-selective (yellows; OTS, LOS, ITG, MTG), and place-selective (greens; CoS, MOG, IPS) regions in the ventral, lateral, and dorsal streams in adolescents ages 10 - 17. *Bottom:* pRF center polar plots same as A but in adults ages 22 - 32. (B) Violin plots of y-position of pRFs in visual degrees for category-selective ROIs in the left hemisphere (light) and right hemisphere (dark) ventral, lateral, and dorsal streams in adolescents (a) and adults (A). ROI colors are the same as in A. Black circle: mean. Error bars: ± SE. Each dot is a participant. (C) Same as B but for eccentricity. (D) Same as B but for pRF size.

### Y position of pRF centers

Building on the observed differences in pRF distributions, we calculated pRF vertical (*y*) position in each participant and ROI (Fig. 3B) and evaluated the stream and category hypotheses as well as any development. Our data reveal no significant differences in vertical pRF positions across age groups (F(1, 51.33) = 1.10, p = 0.30, *LMM: pRF y-position ∼ Age Group x Stream x Category x Hemisphere +(1|Participant);* F(1, 46.43) = 0.27, p = 0.60, *ventral LMM: pRF y-position ∼ Age Group x Category x Hemisphere +(1|Participant)*), nor any interactions between age group and other factors (Supplementary Table 6). These findings suggest that the vertical location of pRFs is mature by adolescence.

Previous studies have largely supported the stream hypothesis, predicting that ventral regions favor the upper visual field and dorsal regions favor the lower visual field^35,36^. However, our results indicate that vertical pRF positions are not determined by stream but instead by category (*LMM*: main effect of category: p = 1.69*10^-22^, F(2, 682.79) = 54.0; *ventral LMM:* main effect of category: F(3,354.26)=54.08; p=8.42*10^-29^; Supplementary Table 6). Across visual cortex, pRFs show category effects as body-selective ROIs display a lower visual field bias (besides left MTG-bodies), place-selective ROIs exhibit an upper visual field bias (besides MOG-places), and face- and word-selective ROIs are located near the horizontal meridian (Fig. 3B). Though category effects are evident, we observed that vertical biases are not always uniform by category but also vary by stream (*LMM*: category x stream effect: p = 6.56*10^-6^, F(2, 682.10) = 12.15), particularly in body- and place-selective regions as noted. Specifically, pRFs in ventral OTS-bodies exhibit a lower visual field bias, but pRFs in lateral MTG-bodies are horizontally biased. Likewise, pRFs in ventral CoS-places have an upper visual field bias whereas pRFs in dorsal MOG-places show a lower visual field bias (Fig. 3A, B).

Unexpectedly, we find significant hemispheric differences in pRF vertical location (*LMM*: p = 1.30*10^-7^, F(1, 686.92) = 28.47; *ventral LMM:* p = 5.66*10^-12^, F(1, 357.27) = 50.81) and significant category by hemisphere interactions (*LMM*: p = 0.03, F(2, 681.69)= 3.44; *ventral LMM:* p = 0.02, F(3, 353.77)= 3.33; Supplementary Table 6). Overall, left hemisphere pRFs are more superior than right hemisphere pRFs. For example, pRFs in left word-selective ROIs are near the horizontal meridian, but pRFs in right word-selective ROIs have a lower visual field bias. These hemispheric differences are pronounced in place-selective regions whereby CoS- and IPS-place ROIs show upper visual field biases in the left hemisphere but equal distribution of pRFs across the upper and lower visual field in the right hemisphere. Together, these data highlight that vertical biases in high-level visual areas mature by adolescence and vary across categories, with additional modulation by stream and hemisphere.

### pRF Eccentricity

With respect to eccentricity, a large body of literature has documented center versus periphery differences in pRF location across category-selective regions, with pRFs in face- and word-selective ROIs exhibiting a central bias – which develops in childhood^27^ – and place-selective regions showing a peripheral bias^22,28,29,32^. We find no significant development in pRF eccentricity from adolescence to adulthood (no significant age group or age group interaction effects: p’s > 0.28, F’s < 1.27; *LMM: pRF eccentricity ∼ Age Group x Stream x Category x Hemisphere +(1|Participant);* p=0.54, F(1,52)=0.27, *ventral LMM: pRF eccentricity ∼ Age Group x Stream x Category x Hemisphere +(1|Participant)*; full stats, Supplementary Table 7).

While pRF eccentricity significantly varies across categories (*LMM:* p=2.12*10^-3^, F(2,685.49) = 6.21; *ventral LMM:* p=2.1*10^-41^, F(3,358.5) = 84.59) with face-selective ROIs displaying more centrally-located pRFs compared to other ROIs (Fig. 3C), our data show that pRF eccentricity is more prominently determined by stream (*LMM:* p = 4.19*10^-19^, F(1,693.18) = 84.62), with pRFs in the lateral stream located, in general, more peripherally than those in the ventral stream (Fig. 3C). This stream effect is further modulated by category (stream x category interaction: p = 5.38*10^-41^, F(2, 684.25)= 106.51; *LMM*) where in the ventral stream, the most peripheral pRFs are in the CoS-places, but in the dorsal-lateral stream, the most peripheral pRFs are in body-selective ROIs (LOS/ITG/MTG; Fig. 3C).

Additionally, there are significant differences in pRF eccentricity across hemispheres (*LMM:* F(1,691.78)=58.60, p=6.5*10^-14^, *ventral LMM:* p=5.53*10^-6^, F(1,364,21)=21.27) whereby pRFs in the left hemisphere are more peripheral than those in the right hemisphere (Fig 3C). Furthermore, we observe significant interactions between stream, category, and hemisphere (*LMM:* p=2.65*10^-5^, F(2,682.72) = 10.7). In the ventral stream, pRFs lie in the central 0° to 5°in right face-selective regions, between 5° to 10° in body- and word-selective ROIs, and extend from 10° to 20° in place-selective regions. But in the dorsal-lateral stream, pRFs in right face- and place-selective regions lie between 5° to 10°, and pRFs in bilateral body- and left face-selective ROIs extend to 10° to 20°. Overall, these data demonstrate that pRF eccentricity is mature by adolescence and that pRF organization in category-selective regions does not segregate simply by category but emerges from the interplay of stream, category, and hemisphere.

### pRF size

While previous studies have not explicitly predicted how pRF size might vary across categories and streams, in retinotopic ROIs pRF size increases linearly with eccentricity. This suggests that pRF size in category-selective regions may mirror pRF eccentricity and exhibit differences across streams, categories, and hemispheres. Additionally, prior work^27^ found that pRF size increases in ventral stream pFus-faces and pOTS-words from childhood to adulthood, but it remains unknown whether pRF size continues to develop during adolescence or in other category-selective ROIs and streams.

Unlike pRF location, pRF size continues to develop from adolescence to adulthood, with significant differences across age group (p=0.02, F(1, 64.2)= 5.62, *LMM: pRF size ∼ Age Group x Stream x Category x Hemisphere +(1|participant)*), age group and hemisphere (F(1,693.04)=9.83, p=1.79*10^-3^), age group, stream, and hemisphere (*LMM:* p=0.04, F(1,692.73) = 4.06), and age group, stream, category, and hemisphere (*LMM:* p=0.03, F(1,683.16) = 3.62; full stats, Supplementary Table 8). Generally, adolescents have larger pRFs than adults, especially in the left hemisphere and in non-face-selective regions (Fig. 3D). Across hemispheres and streams, larger pRF sizes in adolescents compared to adults are more pronounced in the lateral stream than the ventral stream and in body- and place-selective ROIs than face- and word-selective ROIs (besides pSTS-faces, Fig. 3D). In the ventral stream, we do not observe significant development in pRF size (*ventral LMM:* ps>0.05; Fs <3.94; Supplementary Table 8).

Beyond developmental effects, pRF sizes vary significantly by stream (*LMM:* p=3.79*10^-9^, F(1,694.48)= 35.64), with larger pRFs in the dorsal-lateral than ventral stream. PRF size also varies by category (*LMM:* p=7.33*10^-10^, F(2, 686.2)= 21.69; *ventral LMM:* p=1.7*10^-11^, F(3, 359.16.)= 19.07), as pRFs are larger in body- and place-selective ROIs compared to face- and word-selective ROIs. Like pRF position, pRF size shows an interaction between stream and category (*LMM:* p=1.77*10^-11^, F(2,684.82)= 25.68; Supplementary Table 8). For example, in the ventral stream, pRFs in body-selective ROIs are smaller than those in place-selective ROIs while in the dorsal-lateral stream, pRFs in body-selective ROIs are larger than those in place-selective ROIs (Fig. 3D).

Together, these results indicate that pRF size mirrors pRF eccentricity, with regions exhibiting more peripheral pRFs also exhibiting larger pRFs. Furthermore, we find that from adolescence to adulthood, pRF size decreases, particularly in body- and place-selective ROIs and in the dorsal-lateral stream.

### Visual field coverage differentially samples the visual field and continues to mature from adolescence to adulthood

To summarize the collective effect of pRF location and size, we calculated the visual field coverage (VFC) of each ROI (Fig. 4A, B) integrating pRF location and size. VFC captures how pRFs within a region collectively tile the visual field, indicating the total area of the visual field each region processes information from. Given our findings that pRF size continues to develop, decreasing in some regions, we investigated whether this decrease is coupled with smaller VFC in adults compared to adolescents in the corresponding ROIs. Thus, we quantify the full-width half-max (FWHM; Fig. 4A, B - black dotted line) of the VTC in each participant and ROI and compare across age groups, streams, categories, and hemispheres (*LMM: FWHM ∼ Age Group x Stream x Category x Hemisphere +(1|Participant); ventral LMM: FWHM ∼ Age Group x Stream x Category x Hemisphere +(1|Participant)*).

**Figure 4.**
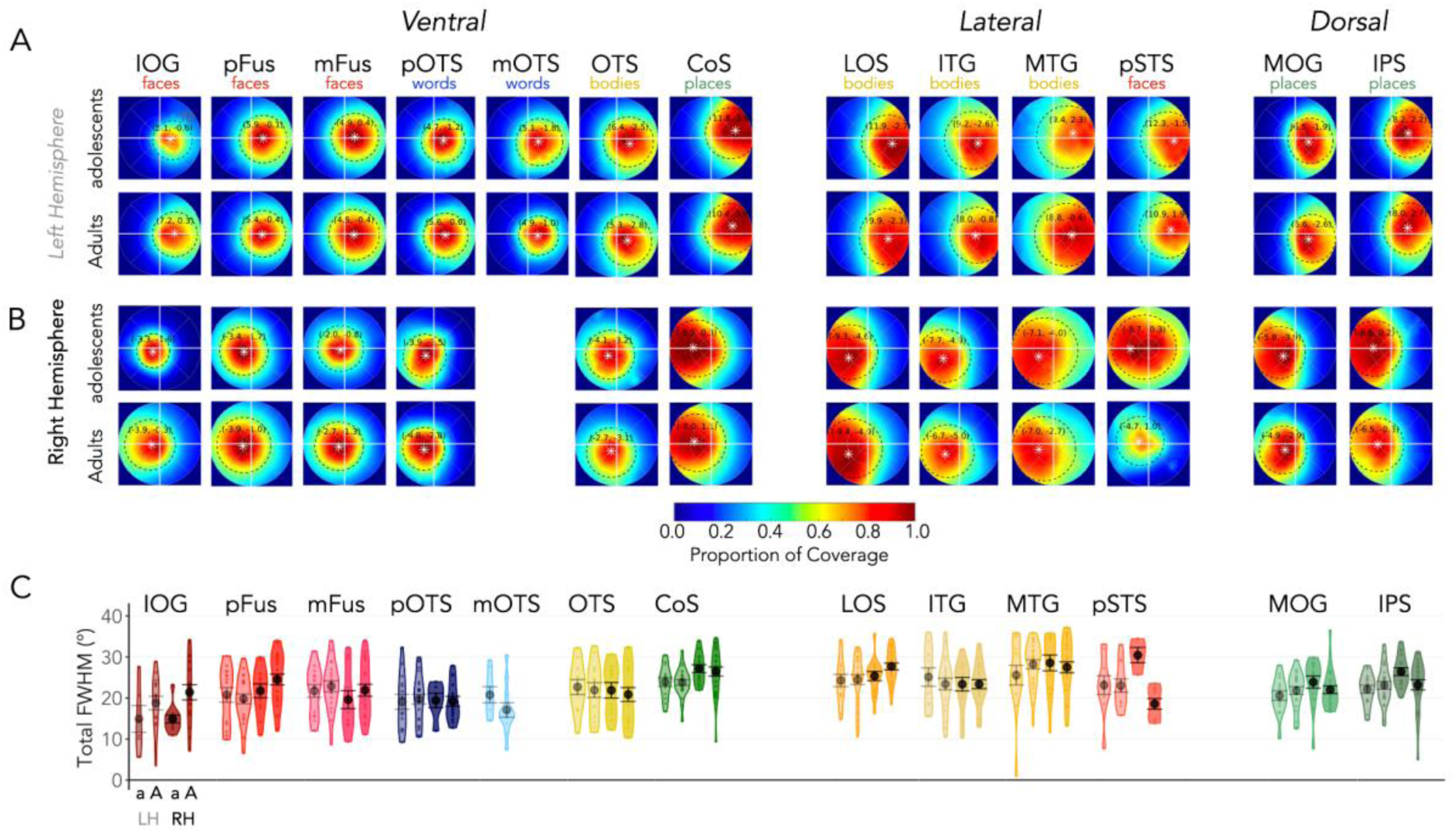
*Visual field coverage (VFC) of category-selective ROIs develops differentially.* (A) Average VFC of each ROI in the left hemisphere across adolescents (top row) and adults (bottom row) hemisphere with a gradient ranging from higher coverage in dark red and lower coverage in dark blue. VFC is calculated as the proportion of pRFs covering each point in the visual field for each participant and is then averaged across participants in the group. *White asterisks with coordinates:* average center of mass of the VFC. Black dotted lines: average full-width half max (FWHM) of the coverage. (B) Same as A but in the right hemisphere. (C) Violin plots of total FWHM in visual degrees for category-selective ROIs in the left hemisphere (light) and right hemisphere (dark) ventral, lateral, and dorsal streams in adolescents (a) and adults (A). Black circle: mean. Error bars: ± SE

In parallel with pRF size, VFC develops differentially from adolescence to adulthood, with total FWHM varying significantly (Fig. 4C) by age group, stream, and category (*LMM:* p = 2.62*10^-3^, F(2, 685.28) = 6) as well as by age group, stream, and hemisphere (*LMM:* p = 7.68*10^-3^, F(1, 693.29) = 7.15; Supplementary Table 9). VFC development is more pronounced in the right hemisphere, in the dorsal-lateral stream, and in face ROIs, as we observe significant decreases in overall coverage of the visual field for right pSTS-faces (post-hoc t-test: t(27.06) = 5.20, p = 1.98*10^-3^) and right IPS-places (post-hoc t-test: t(37.99) = 2.06, p = 0.05) and significant increases in coverage for right IOG-faces (post-hoc t-test: t(6.04) = -3.09, p = 4.61*10^-3^) from adolescence to adulthood.

In addition, we observe differences in VFC across streams (*LMM:* p = 0.05, F(1,698.78) = 4), with the ventral stream displaying a central bias (∼30% of the FWHM in the center 5° of the visual field) and the dorsal-lateral stream showing more peripheral coverage (just ∼15% of the FWHM in the central 5°, Fig. 4A, B; Supplementary Table 10). Additionally, the lateral ROIs exhibit more contralateral VFC than both ventral stream ROIs and dorsal place ROIs (Fig. 4A,B; Supplementary Table 10). VFC also differed significantly by category (*LMM:* p = 0.02, F(2,689.52) = 3.85; *ventral LMM:* p = 1.55*10^-8^, F(3, 359.39) = 13.80; Supplementary Table 9). For example, the ipsilateral visual field coverage is larger for face-selective ROIs than place and body ROIs (Fig. 4; Supplementary Table 10). Notably, like with pRF location and size, VFC varied by stream and category (*LMM:* p = 9.58 * 10^-7^-, F(2,689.01) = 14.14; Fig. 4C), as there is a central bias in ventral face- and body-selective ROIs (∼35% of FWHM in central 5°) but a peripheral bias in their lateral counterparts (∼17% of FWHM in the central 5°, Supplementary Table 10). Place-selective ROIs, however, had similarly peripheral coverage of the visual field across streams (∼13%, Fig. 4A, B), even as their upper vs. lower visual coverage varied across streams. Overall, we find the largest central bias in the right ventral face ROIs, in which over 50% of the VFC is concentrated in the central 5°.

Collectively, analysis of VFC reveals both decreases and increases in VFC of category-selective ROIs from adolescence to adulthood.

### Category selectivity continues to develop during adolescence alongside spatial computations

Our findings reveal that while pRF location is mature by adolescence, pRF size and VFC continue to develop into adulthood, raising important questions about how these changes relate to the development of category selectivity. As previous work finds that category selectivity continues to mature into adolescence^10,37–39^, we examined if category responses develop in these same participants and ROIs and whether this development might be linked with the development of pRFs.

To do so, we quantified category selectivity (mean t-value) in 10mm disks ROI centered on each category ROI and tested if selectivity varied across age group, stream, category, and hemisphere (Fig. 5A; *LMM: mean t-value ∼ Age Group x Stream x Category x Hemisphere +(1|Participant); ventral LMM: mean t-value ∼ Age Group x Category x Hemisphere +(1|Participant)*). We find differential development of category selectivity from adolescence to adulthood (significant age group by category interactions, *LMM:* p = 2.36 * 10^-7,^ F(2, 746.92) = 15.58; *ventral LMM:* p = 1.81*10^-3^, F(3, 358.77) = 5.1; Supplementary Table 11). This development is associated with significant decreases in place selectivity across all place ROIs, bilaterally, (post-hoc t-tests: p’s <0.01, t’s > 2.39), and significant increases in face selectivity in right hemisphere IOG-faces (post-hoc t-tests: p = 0.02, t(28.27) = -2.40; Fig. 5A). Additionally, we find that the size of these regions also develops into adolescence (Supplementary Table 12).

**Figure 5.**
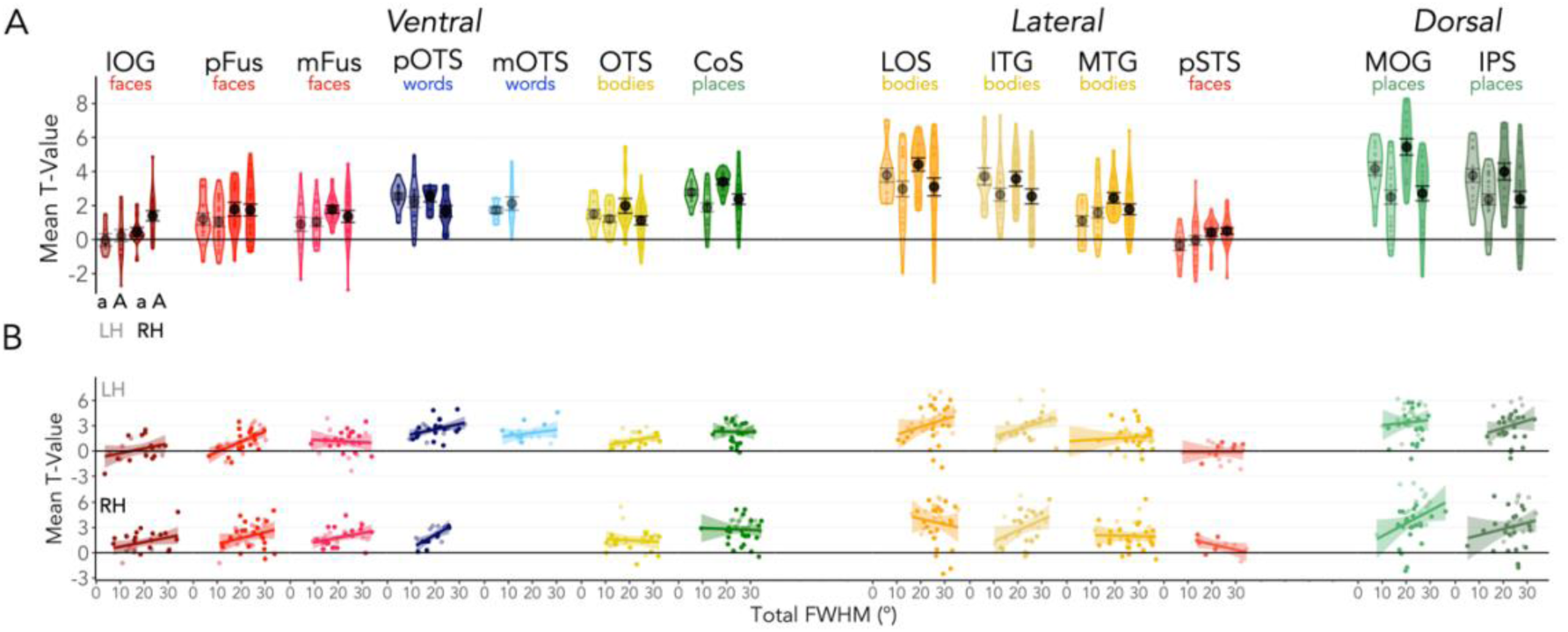
*Category-selectivity differentially develops from adolescence to adulthood. (*A) Violin plots of average t-values for the category contrast for the preferred category of each ROI (e.g., faces > all other categories for the IOG-face ROI) in the left hemisphere (light) and right hemisphere (dark) ventral, lateral, and dorsal stream ROIs in adolescents (a) and adults (A). Black circle: mean. Error bars: ± SE. B. Linear relationships between total FWHM and mean t-value in each ROI in the left hemisphere (top) and right hemisphere (bottom). Each dot is a participant; adolescents are colored in lighter colors.

Given that VFC and category selectivity both develop and are each related to visual behaviors like reading and face recognition^24,37,44,45^, we examined if these properties are linked. We find a significant link between category selectivity and VFC (Fig. 5B; p = 0.03, F(1, 628.47) = 4.67, *LMM: mean t-value ∼ total FWHM x Stream x Category x Hemisphere +(1|Participant)*; p = 9.45*10^-5^, F(1, 323.53) = 15.63, *ventral LMM: mean t-value ∼ total FWHM x Category x Hemisphere +(1|Participant)*; full stats, Supplementary Table 13), which additionally varies by stream and category (significant interaction FWHM x category x stream: p = 6.81*10^-3^, F(2, 624.3) = 5.03, *LMM*). That is, in left pFus-faces, LOS-bodies, and IPS-places, as well as right pOTS-words, ITG-bodies, and MOG-places (but not other ROIs) higher category selectivity is linked with greater VFC (Fig. 5B).

Together, these results suggest that category selectivity continues to develop after pRF location has matured and parallels the development of VFC with larger category selectivity associated with larger VFC.

## Discussion

Here, we examined pRFs in category-selective regions across the ventral, dorsal, and lateral visual streams and chart their development from adolescence to adulthood using a novel experiment that drives high-level regions. Across all ages, we find the location and sizes of individual pRFs, as well as their combined VFC, vary systematically across both category and stream. While the location of pRFs remains stable from adolescence to adulthood, we find that pRF size and VFC continue to develop differentially during adolescence alongside category selectivity. Functionally, our findings suggest that visuospatial processing is not governed by a single principle but by an interplay of category and stream, as well as hemisphere, necessitating a rethinking of visuospatial computations by pRFs in visual cortex. Developmentally, we find that the area of the visual field in which visual information is integrated shows a protracted refinement through adolescence, underscoring the importance of studying the development of the adolescent brain.

### Visuospatial computations in high-level visual cortex vary by both stream and category

Prior research has implicated either category or stream in how category-selective regions sample the visual field. Eccentricity Bias Theory^29^ highlights the relation between category selectivity and eccentricity based on evidence that ventral face- and word-selective regions have central biases, whereas place-selective regions have peripheral biases ^22,27–29,31,32,46^. Stream Theory^35,36^ highlights the coupling between stream and vertical biases, suggesting that like biases in EVC, ventral stream regions have an upper visual field bias and dorsal stream regions have a lower field bias. However, our results suggest that pRF eccentricity, vertical position, size, and VFC vary across streams, categories, and hemispheres. Surprisingly, pRF eccentricity varies more by stream than by category, with ventral stream pRFs located more centrally than dorsal and lateral streams pRFs which extend to the periphery, while pRF vertical bias varies more by category than by stream, with body-selective regions exhibiting more pRFs in the lower visual field, place-selective regions having more pRFs in the upper visual field, and pRFs in face- and word-selective regions more concentrated along the horizontal meridian. Importantly, we find that the contributions of category and stream for pRF eccentricity and vertical position are not mutually exclusive but rather exhibit significant interactions: pRFs in ventral face and body-selective regions are smaller and more central than those in the place selective regions, but the pattern is reversed in dorso-lateral regions. Finally, pRFs and VFC in right hemisphere category-selective regions extend more into the ipsilateral and lower visual field compared to the left hemisphere. These findings challenge previous hypotheses and suggest that different dimensions than previously posited drive differences in pRF eccentricity and vertical position in high-level visual regions.

We hypothesize that retinotopic regularities associated with viewing specific categories in the context of different tasks may shape the differential pRFs properties across category-selective regions, streams, and hemispheres. Prior research has theorized that retinotopic biases in category selective regions are tied to the regularity of different categories in specific locations in the visual field^22,24,27–30,32,47–50^. For example, ventral regions associated with categories that observers tend to fixate upon – faces, words, and even cultural objects like Pokemon^48,51–54^ – have a foveal bias^28,32,55,56^. Interestingly, the tendency to fixate on faces produces higher occurrences of bodies and limbs in the lower visual field^30,51^, reflected in the lower field bias of body-selective regions^30^. In the real world, places extend to the periphery, reflected in the peripheral bias of place-selective regions^22,28,32,57^. We hypothesize that differential pRFs across categories and streams may be a result of the statistical regularity of visuospatial information during specific tasks. That is, pRFs and population codes across pRFs spanning a region may be optimized for processing category-relevant and task-specific information.

Indeed, several studies have shown a link between the population code spanned by pRFs tiling a region and behavior^12,23,24,27,30,47,58,59^. For instance, spatial computations in TOS-places predict navigational affordances^47^ and differences in pRFs for upright versus upside-down faces in ventral face-selective regions predict the face inversion effect^24^. Thus, we hypothesize that visual navigational affordances, a task associated with TOS-places in the dorsal stream^60–62^, are more prominent in the lower-visual field and require more central processing than recognition of places^29,63,64^, but features in the periphery as well as in the upper visual field (e.g., sky, ceiling) may aid scene classification supported by CoS-places. Likewise, processing of visual dynamics and social information associated with body and face regions in the lateral stream^2–4,6^ may require processing of biological motion in the periphery, whereas face recognition^65^ and learning to read^40^ (and write^66^) tasks associated with the ventral stream, may be associated with fixation on faces, words, and hands, respectively. This hypothesis can be tested with advances in mobile eye tracking that can be used to quantify observers’ visual diet (occurrence of different categories in different parts of the visual field) under different tasks.

### pRF location is mature by adolescence but spatial integration continues to develop

Across adolescents and adults, we find no quantitative or qualitative differences in pRF eccentricity or vertical position in category-selective regions in all visual streams. These data suggest that pRF location is mature by adolescence, consistent with prior work showing that pRFs in EVC (V1-V3) are mature by childhood^27,41,42^.

In contrast to pRF location, we find that pRF size and VFC continue to change during adolescence, as does category selectivity in the same individuals. Previous work^27^ finds that pRFs in ventral face and word-selective regions develop during childhood. Our results suggest that the area of the visual field from which neurons in high-level face- and place-selective regions spatially integrate information also continues to develop during adolescence, and the differential development across hemispheres may be associated with functional specialization^55,67^. Furthermore, in addition to corroborating the hypothesis that category selectivity continues to develop into adolescence^10,37–39^, our findings reveal a nuanced timeline of visual development, illustrating that pRF location matures first and may support the ongoing maturation of pRF size, VFC, and category selectivity – which develop in parallel.

The dynamic nature of visual development is exemplified by the differential changes observed in category selectivity and VFC. VFC and category selectivity increase for right hemisphere IOG-faces but decrease for right hemisphere place-selective regions highlighting the coupling between functional specialization and the underlying spatial computations. Critically, this sheds light on the fact that cortical development is not just associated with expansion but also with reduction. This pattern of development has important implications for theories regarding how cortex is recycled to adapt for changing behaviors over development^37,56^. Behaviorally, mature pRF location in adolescents’ high-level visual areas across streams suggest the possibility that adolescents have similar fixation patterns as adults under different tasks. Nonetheless, developmental changes in pRF size and VFC suggest differential spatial integration across adolescents and adults, which may correlate with differential performance in specific behaviors, e.g., in face recognition, reading, and navigation^12,24,27,47,58^.

It is interesting to consider the mechanisms underlying these differential developments. Prior research suggests that eccentricity biases in high-level visual cortex mirror white matter connections^22,68^ and functional connectivity^69–72^ to eccentricity bands in EVC. For example, ventral face- and word-selective regions have more white matter connections to EVC in the central 5° than in peripheral bands, while place-selective regions have more connections to peripheral EVC bands than the central 5°^22,68^, and these connectivity patterns are present at birth^68^. Together, these discoveries suggest that earlier development of pRF location may be established by long range white matter connections that are present in infancy. However, the distribution of white matter connections between category-selective regions in the lateral and dorsal stream to EVC as well as their development is unknown, and can be tested in future research. We hypothesize that the later maturation of pRF size may be related to protracted development of dendritic arborization and synaptic weights in high-level visual cortex^73,74^, which may affect spatial pooling. Another possibility is that changing visual behaviors during adolescence might affect the area over which information is spatially integrated. Future research examining the longitudinal development of adolescents’ visual diet, together with longitudinal measurements of pRFs and category selectivity, may shed light on how one’s visual diet and behaviors impact spatial integration and category selectivity.

Overall, our study suggests a rethinking of spatial computations in high-level visual cortex and their development during adolescence. As such, our research sets a new foundation for future investigations into how visual experience and viewing patterns may sculpt visuospatial computations across development and how this processing might diverge in cases of atypical development like autism^75^, dyslexia^40,76^, or prosopagnosia^77–79^ where altered viewing patterns may also alter visuospatial computations.

## Methods

### Participants

19 neurologically typical adolescents aged 10 to 17 years old (M = 13.74 ± 2.13; 11 females, 8 males) and 27 adults (ages 22 - 32; M = 25.52 ± 3.00; 13 females, 14 males) participated in this study. 4 of the adolescents participated in two scanning sessions approximately a year apart. All participants had normal or corrected-to-normal vision. Participants, or their parents, gave written informed consent, and all procedures were approved by the Stanford Internal Review Board on Human Subjects Research.

Sessions were excluded from the analysis if within scan motion and/or between scan motion was greater than 2 voxels. Of the 19 adolescent sessions, 4 participants were excluded based on these motion criteria. No adult sessions were excluded. After exclusion, we analyzed the data of 15 adolescents (ages 10 - 17; 9 females, 6 males) and 27 adults (ages 22 - 32; 13 females, 14 males).

### Data Acquisition

#### MRI

Participants were scanned using a General Electric Discovery MR750 3T scanner located in the Center for Cognitive and Neurobiological Imaging (CNI) at Stanford University. A phase-array 32-channel head coil was used for the category localizer experiment and to obtain anatomical scans. For the toonotopy experiment, a 16 channel head coil was used.

#### Anatomical scans

For each participant, we obtained a whole-brain anatomical volume using a T1-weighted pulse sequence (TI = 450ms, 1x1x1mm, flip angle = 12 degrees, FoV = 204 mm). Anatomical images of each brain were used for segmentation of the gray/white matter boundary.

#### Toonotopy

Participants completed four runs of a wide-field pRF mapping fMRI experiment with cartoon stimuli, which we refer to as Toonotopy^22^. In the experiment (Fig. 1A), bars of width 5.7° swept a circular 40°x40° (visual angle) aperture with a fixation dot at center.

The bars swept the visual field at four orientations (0°, 45°, 90°, 135°) in eight directions (2 opposite directions orthogonal to each orientation). The cartoon stimuli randomly changed at a rate of 8 Hz with blanks (mean luminance gray background with fixation) appearing at regular intervals. During each run, participants fixated on the central dot and were instructed to press a button whenever the dot changed colors. Each run was 3 minutes and 24 seconds long.

#### Category localizer

The same participants underwent an fMRI category localizer experiment, which is used to identify voxels whose neural response is stronger to one category vs. many other categories^43^. In each run, participants were presented with stimuli from five domains, each with two categories (Fig. 2A; faces: child, adult; bodies: whole, limbs; places: corridors, houses; objects: cars, guitars; characters: words, numbers). Images within the same category were presented in 4s blocks at a rate of 2Hz and were not repeated across blocks or runs. 4s blank trials were also presented throughout a block. During a run, each category was presented eight times in counterbalanced order, with the order differing for each run. Throughout the experiment, participants fixated on a central dot and performed an oddball detection task, pressing a button when phase-scrambled images randomly appeared. Each participant completed 3 runs of the category localizer experiments with different images; each run was 5 minutes and 18 seconds long.

### Data Analysis

#### Anatomical data analysis

T1-weighted images were automatically segmented using FreeSurfer (FreeSurfer 7.0.0:^80^) and then manually validated and corrected using ITKGray. Cortical reconstructions were generated from these segmentations using FreeSurfer.

#### fMRI data analysis

Data were processed and analyzed in MATLAB using mrVista (http://github.com/vistalab). All data were analyzed within the individual participant native brain space. Functional data were manually aligned to the T1-weighted volume. The manual alignment was then optimized using robust multiresolution alignment. For participants with more than one session, functional data were aligned to the anatomical scan taken closest to the date of the functional scan. Data were not spatially smoothed and were restricted to the cortical ribbon. Functional data were motion-corrected within and between scans using mrVista motion correction algorithms. Quality assurance was also performed to determine exclusions based on motion.

#### mrVista to FreeSurfer conversion

To visualize functional maps and draw regions of interest (ROIs), eccentricity and phase maps from Toonotopy and category selectivity maps from the category localizer experiment were converted from mrVista to FreeSurfer coordinates and projected onto the inflated cortical surface reconstruction for each individual participant in Freeview (FreeSurfer 7.2.0).

#### Defining ROIs

ROIs were drawn using Freeview (FreeSurfer 7.2.0) on the inflated cortical surface reconstruction of each participant’s brain then projected back to mrVista for analysis. ROIs were defined by JKY and JOC.

*V1-V3:* Using polar angle and eccentricity maps from the Toonotopy experiment thresholded at 20% variance explained, we defined early retinotopic visual areas (V1, V2, V3; Fig. 1B) in both hemispheres in each individual. Boundaries between retinotopic areas were defined as the middle of polar angle reversals at the horizontal or vertical meridian representations, and each area included foveal to peripheral representations^14,81^. Dorsal (V1d, V2d, V3d) and ventral (V1v, V2v, V3v) components of each visual area were defined separately and combined in analysis to create representations of the entire visual field (V1, V2, V3).

*Category ROIs:* From the category localizer experiment, statistical contrast maps of each category domain versus all other category domains (i.e. faces > all other stimuli) were thresholded at a *t-value* > 3 at the voxel level, as in previous experiments^22,27,43^. Using these contrast maps, category-selective ROIs in the ventral, lateral, and dorsal streams were defined in each subject as clusters of voxels selective for a category located at a particular anatomical landmark (Fig. 2B).

Face-selective voxels (contrast: adult and child faces > all other categories) were defined in the inferior occipital gyrus (IOG-faces), posterior fusiform gyrus (pFus-faces), mid fusiform gyrus (mFus-faces), and posterior superior temporal sulcus (pSTS-faces). Word-selective voxels (contrast: words > all other categories except numbers) were defined in the posterior occipital temporal sulcus (pOTS-words) and mid occipital temporal sulcus (mOTS-words). Because we could identify only in a minority of subjects (∼20%) the right mOTS-words, mOTS-words was only defined in the left hemisphere. Bodypart-selective voxels (contrast: bodies and limbs > all other categories) were defined in the occipital temporal sulcus (OTS-bodies), lateral occipital sulcus (LOS-bodies), inferior occipital gyrus (IOG-bodies), and mid temporal gyrus (MTG-bodies). Place-selective voxels (contrast: corridors and houses > all other categories) were defined in the collateral sulcus (CoS-places), intraparietal sulcus (IPS-places), and mid occipital gyrus (MOG-places^82,83^).

Because we were only able to identify pSTS-face and MTG-bodies in a minority of adolescents (pSTS-faces: left 0/15; right 3/15; MTG-bodies: left: 5/15, right 4/15) we used group maximum probability maps (MPM) of these ROIs from adults projected to individual cortical surfaces for both adolescents and adults. The MPMs were thresholded at voxels found in 30% or more of the participants. After defining the category ROIs, only regions with ten or more voxels and more than 20% variance explained during the Toonotopy experiment were included in subsequent analyses (see Supplementary Table 2 for number of subjects per ROI).

Overall, we report data from 7 category-selective ROIs in the ventral stream (IOG-faces, pFus-faces, mFus-faces, pOTS-words, mOTS-words, OTS-bodies, CoS-places), 4 ROIs in the lateral stream (pSTS-faces, LOS-bodies, ITG-bodies, MTG-bodies), and 2 ROIs in the dorsal stream (MOG-places, IPS-places). As prior research combined the dorsal and lateral stream into a single dorsal stream^25,32,36^ and because streams differ in the number and type of category-selective regions – e.g., dorsal stream contains only one category (places) – throughout this study we compare the ventral stream and a combined dorsal-lateral stream.

#### Estimating pRFs

The time-course data were transformed from functional slices to the T1-weighted whole brain anatomy using trilinear interpolation. The pRF of each voxel was modeled using the compressive spatial summation (CSS) model^19^ (Fig. 2B) using VISTA lab software (http://github.com/vistalab). A pRF is modeled independently for every voxel by fitting a 2D Gaussian with a center (x,y) and a size determined by the standard deviation (σ) of the Gaussian in degrees of visual angle. An exponent parameter (0≤n≤ *1*) is additionally fit for each voxel to capture the compressive nonlinearity of pRFs. pRF size is thus defined as 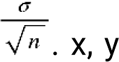, and σ are iteratively optimized to minimize the root mean squared error between the observed and predicted time-series. Eccentricity 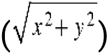 and phase 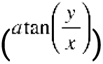 of each voxel were derived from the center (x,y) coordinates and used to generate eccentricity and phase maps, respectively. As in prior studies^22,24^ we report pRF parameters for voxels in which the variance explained by the pRF model is greater than 20%.

*pRF size vs. eccentricity.* To evaluate the relationship between pRF size and eccentricity (Fig. 1), all analyzed voxels in each participant’s ROI were entered into a linear regression comparing pRF size to eccentricity. A line of best fit was derived for each participant, and the slope and intercept of the line was averaged across participants of each group (Fig. 1D).

#### Visual field coverage (VFC)

VFC, the region of the visual field processed by the set of pRFs spanning an ROI, was calculated for each ROI and participant. RFs were represented by a binary circular mask centered on their centers (x,y) with size σ; coverage was calculated by determining the density of pRFs at each point. To create group VFC maps (Fig. 4A, B), we averaged the individual VFC maps, whereby each visual field location illustrates the average pRF density, for that ROI per each group (adolescents, adults).

##### Estimating the full-width half-max (FWHM) of the VFC

For each ROI and participant, we calculated the FWHM (Fig. 4A, B - black dashed line), which provides a standardized measure of the spatial extent of the VFC by estimating the diameter, in visual degrees, of the cross section of the VFC in which it reaches half of its maximum amplitude. The FWHM was determined by fitting a circular Gaussian centered at the center of mass, namely, the peak response (Fig. 4A, B - white asterisk), of the VFC, using the equation: 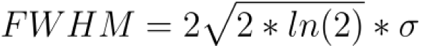 where *σ* is the standard deviation of the Gaussian.

##### Proportion of FWHM measures

We quantified the proportion of the FWHM in (i) each of the four quadrants of the visual field: upper contralateral (UC), lower contralateral (LC), lower ipsilateral (LI), and upper ipsilateral (UI) quadrants of the visual field and (ii) separately in the central 5°. Estimates were done separately for each ROI, hemisphere, and participant. This approach provides a detailed analysis of VFC across different field quadrants and eccentricity (Supplementary Table 10).

### Category selectivity analysis

To examine the development of category selectivity in our functional category ROIs, we quantified the mean t-value and the ROI size for each ROI in each individual and then compared across age groups.

#### Mean t-value

To evaluate how the strength of category selectivity develops, we quantified the mean t-value, which indicates how strongly an ROI responds to one visual category versus all other categories. To control for ROI size differences across participants, groups, and ROIs and focus on selectivity, we generated a 10mm radius disk ROI centered on the original functional category ROI. The 10mm disk ROI approximated the average ROI size across all participants and regions. We then calculated the mean t-value (unthresholded) of the 10mm disk ROI for the category and ROI was drawn for (Supplementary Table 11).

#### ROI size

To examine the extent of cortex involved in processing each category and how this develops from adolescence into adulthood, we also measured ROI size. Using our original category ROIs, we quantified the number of category-selective voxels with t-value > 3 in each category ROI for every participant (Supplementary Table 12).

### Statistical Analysis

All statistical analyses were conducted using R version 4.2.2. All error bars in the main and supplementary figures represent the standard error of the mean across participants in a group. Except for analysis of EVC which included all subjects, the number of participants included in each statistical test, based on whether or not they had an ROI, was consistent across all analyses and can be found in Supplementary Table 3.

We use two sets of linear mixed effect models (LMMs, two-tailed) analyses to quantify significant differences across age group, stream, category, and hemisphere. In the first analysis, referred to as LMM, we compared age groups, streams, categories, and hemispheres for face, body, and place ROIs, which are distributed across multiple streams. As prior research combines the dorsal and lateral stream into a single dorsal stream and because streams differ in the number and type of category-selective regions – e.g., dorsal stream contains only one category (places) – we compare the ventral stream and a combined dorsal-lateral stream. In the second analysis, referred to as ventral LMM, we assessed differences across age group, category, and hemisphere for ROIs within the ventral stream in order to include word-selective ROIs, which were only found in the ventral stream.

#### LMMs

were conducted using the lme4^84^ and emmeans packages in R (https://CRAN.R-project.org/package=lme4, https://CRAN.R-project.org/package=emmeans). Each dependent variable, including pRF parameters (x-position, y-position, size, eccentricity), visual field coverage (FWHM), and category selectivity (mean t-value, ROI size), was analyzed using a series of LMMs to account for both fixed and random effects, repeated measures, as well as incomplete data (e.g., not all participants had all ROIs). The primary fixed effects included age group (adolescents, adults), stream (ventral, dorsal-lateral), category (faces, words, bodies, places), and hemisphere (left, right), with participant as a random effect to control for inter-subject variability. We additionally conducted analyses with age as a continuous variable for comparison (Supplementary Tables 3–11). Given the variability in the number and type of category-selective regions found across streams – e.g., word-selective ROIs were exclusively found in the ventral stream, and place-selective ROIs dominate the dorsal stream – analyses were structured in two complementary approaches:

First, we assess differences across streams, categories, groups, and hemispheres across face, body, and place ROIs which are found in multiple streams (1; referred to as LMM). For this analysis, we excluded word ROIs and combined the dorsal and lateral stream into a single dorsal-lateral stream, as prior studies did not separate these streams, and this enables comparisons between categories shared across the dorsal-lateral and ventral streams.

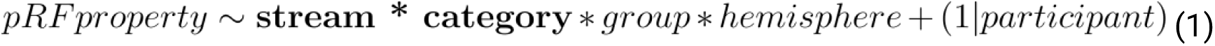

Second, we assess differences across age group, category, and hemisphere for ROIs within the ventral stream (2; referred to as ventral LMM). This analysis allows us to compare word-selective ROIs, which were only found in the ventral stream.

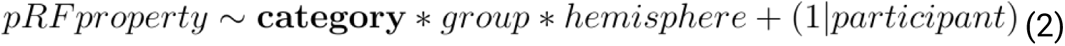

Full results and statistics from all LMMs and ventral LMMs are included in Supplementary Tables 1 - 13. Additional analyses with age as a continuous variable, rather than by group, is also included in the Supplementary Tables 1 - 13.

#### Post-hoc testing

Post-hoc unpaired, independent t-tests (two-tailed) were performed using the t.test function in R to further examine significant main effects of age group and age group interactions from the LMMs. For each ROI, t-tests compared the mean value of the dependent variable between adolescents and adults. Post-hoc t-tests were conducted for pRF size, total FWHM, category selectivity (mean t-value), and ROI size, given main effects of or interactions with age group. Only significant post-hoc t-tests are reported in the main text.

### Author Contributions

J.K.Y. analyzed the data and wrote the manuscript. J.C. contributed to data analysis and contributed to the manuscript. D.F. designed the experiment, collected data, and contributed to the manuscript. K.G-S. designed the experiment, contributed to data analysis, and wrote the manuscript.

## Acknowledgements

This material is based upon work supported by the National Science Foundation Graduate Research Fellowship Program under Grant No. (DGE-2146755) awarded to J.K.Y., NIH grants (grant numbers RO1EY022318, RO1EY023915 to K.G.S.), a William R. and Sara Hart Kimball Stanford Graduate Fellowship awarded to D.F., and a Symbolic Systems Summer Internship Program internship (J.C.).

## Competing Interests

The authors declare no competing interests.

## Data Availability

Raw data available upon request to J.K.Y. Processed data available on Github: https://github.com/VPNL/toonCat

## Code Availability

fMRI data were analyzed using the open source mrVista software package (https://github.com/vistalab/vistasoft). Custom code for processing the pRF experiment and functional localizer, reproducing figures, and statistics can be found at https://github.com/VPNL/toonCat.

## Additional Information

## Supplementary information

See attached.

**Correspondence and requests for materials** should be addressed to J.K.Y.

## Notes

https://github.com/VPNL/toonCat

### Competing Interest Statement

The authors have declared no competing interest.

