## Supplementary Figures and Tables for "Visuospatial computations vary by category and stream and continue to develop in adolescence"

**Supplementary Figure 1: Phase maps for all participants.**

Phase maps for each individual participant thresholded at 20% variance explained on left and right hemisphere medial inflated surfaces. All analyses were done within each participant's brain. Each pair of panels (left, right hemisphere) shows one participant, organized from youngest (left) to oldest (right) within each row. Age indicated in black at the top left corner of each pair of hemispheres. Color wheel on bottom right indicates pRF phase in visual degrees.

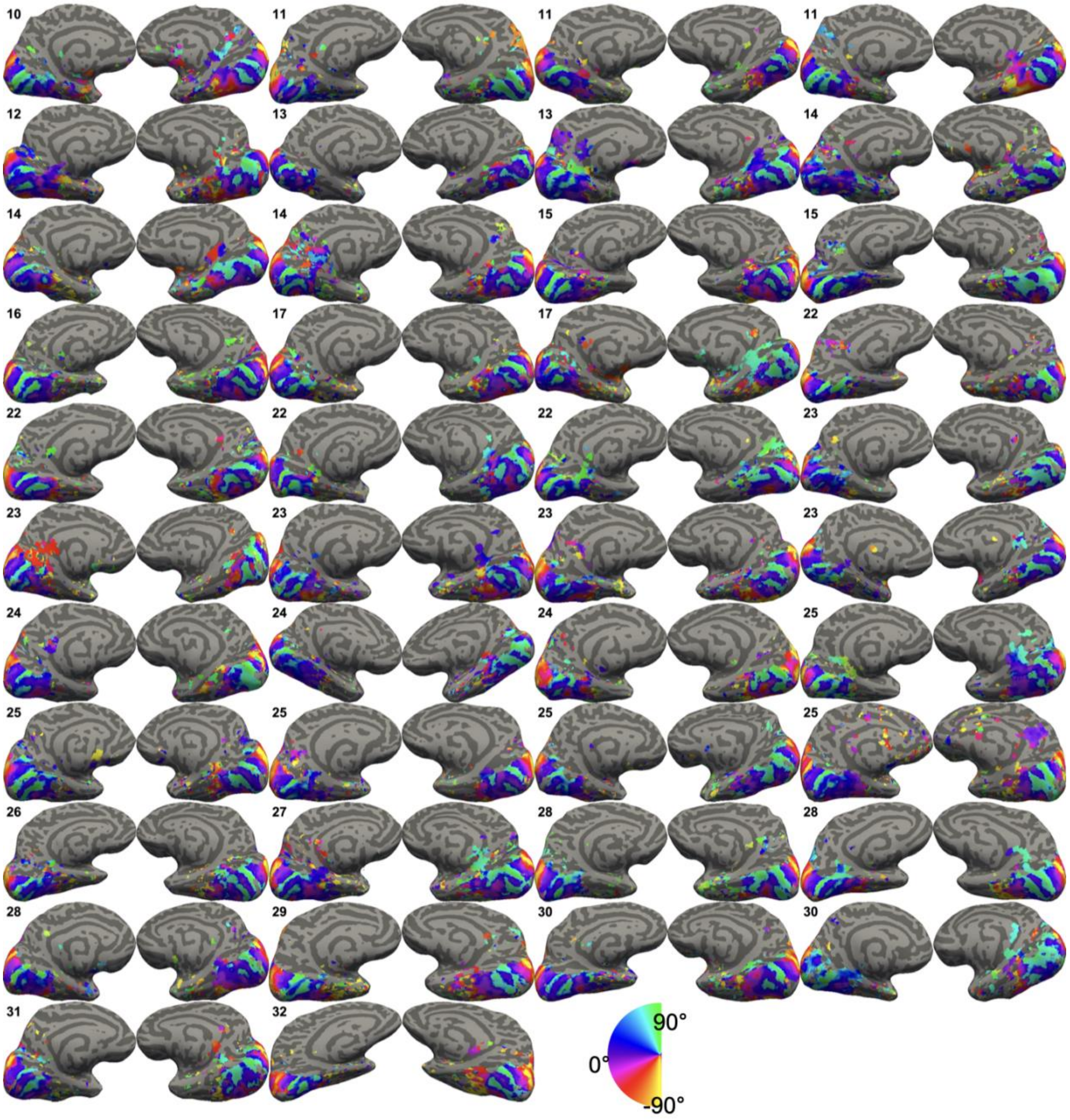

**Supplementary Figure 2. Eccentricity maps for all participants.**

Eccentricity maps for each individual participant thresholded at 20% variance explained on left and right hemisphere medial inflated surfaces. All analyses were done within each participant's brain. Each pair of panels (left, right hemisphere) shows one participant, organized from youngest (left) to oldest (right) within each row. Age indicated in black at the top left corner of each pair of hemispheres. Color wheel indicates pRF eccentricity in visual degrees.

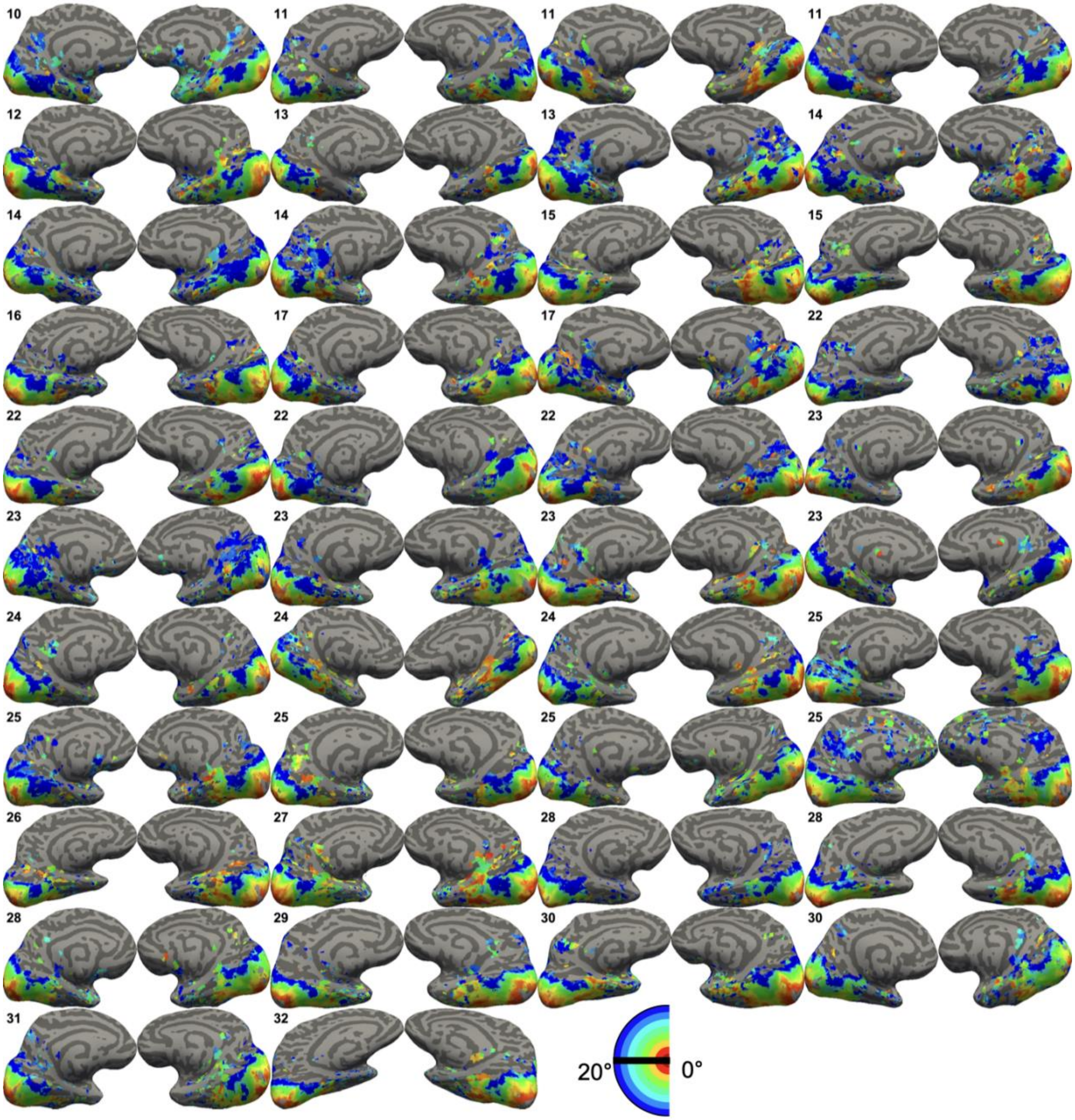

**Supplementary Figure 3. Size maps for all participants.**

Size maps for each individual participant thresholded at 20% variance explained on left and right hemisphere medial inflated surfaces. All analyses were done within each participant's brain. Each pair of panels (left, right hemisphere) shows one participant, organized from youngest (left) to oldest (right) within each row. Age indicated in black at the top left corner of each pair of hemispheres. Color wheel indicates pRF size in visual degrees.

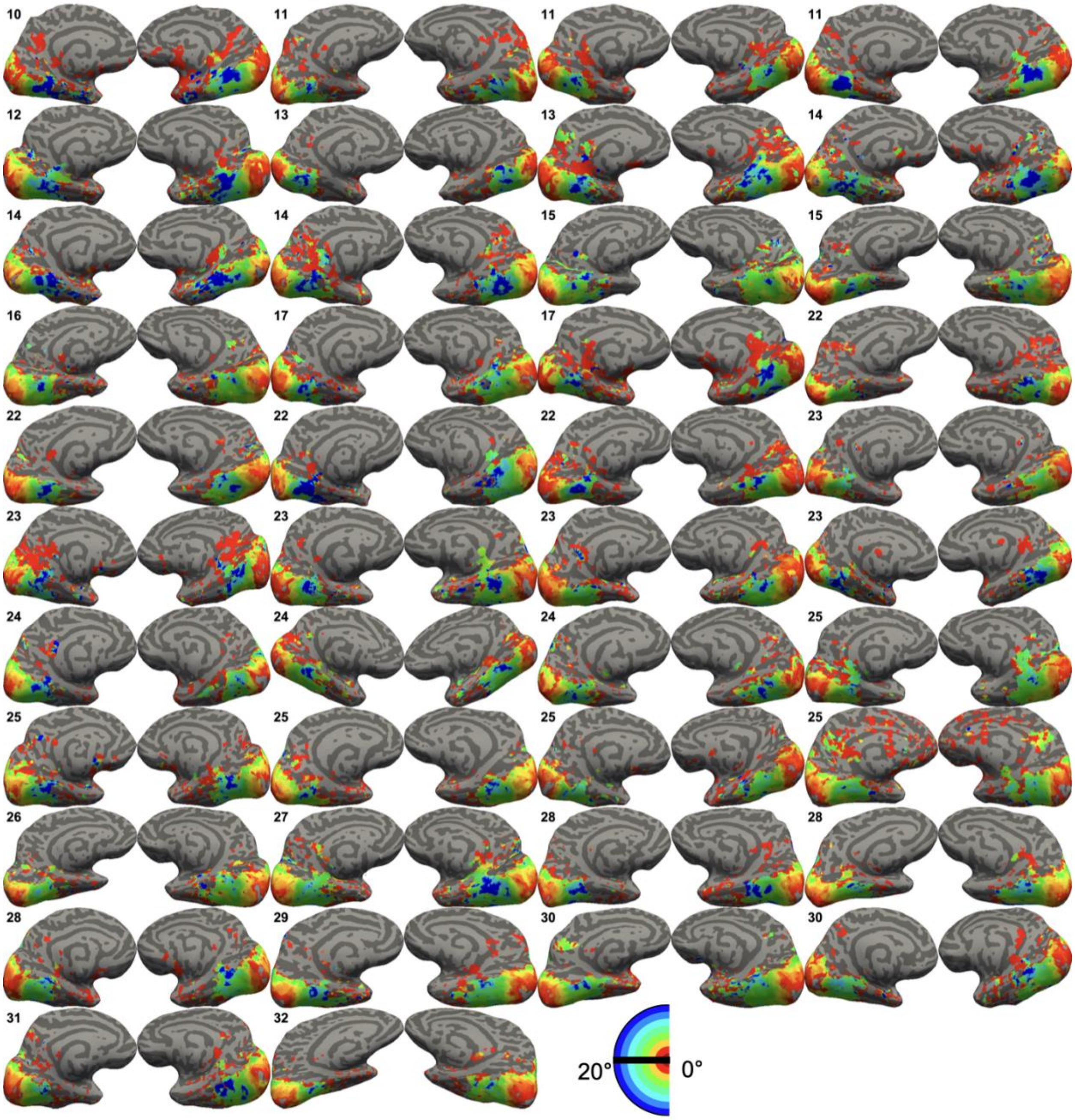

**Supplementary Figure 4: pRF x-position.**

Violin plots of y-position of pRFs in visual degrees for category-selective ROIs in the left hemisphere (light) and right hemisphere (dark) face-selective (reds; IOG, pFus, mFus, pSTS), word-selective (blues; pOTS, mOTS), body-selective (yellows; OTS, LOS, ITG, MTG), and place-selective (greens; CoS, MOG, IPS) regions in the ventral, lateral, and dorsal streams in adolescents (a) and adults (A). Error bars: ± SE (standard error of the mean).

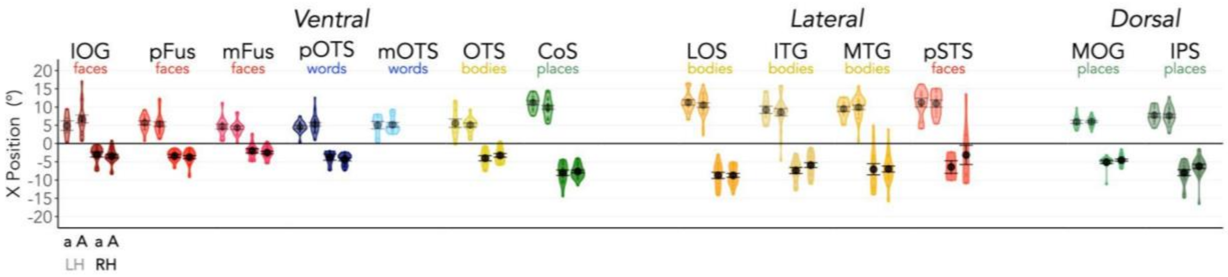

**Supplementary Table 1: Differences in pRF parameters across early visual cortex.** Results of LMMs (*parameter ~ Age Group x ROI (V1/V2/V3) x Hemisphere + (1| participant)*) examining differences across factors: age group, ROI, and hemisphere in early visual cortex (V1, V2, V3) for pRF center x-position, pRF center y-position, pRF eccentricity, and pRF size. We find no significant effect of age group across all parameters in early visual cortex.

| Analysis | model term | df1 | df2 | F.ratio | p.value |
|---|---|---|---|---|---|
| LMM: pRF centers X-position (Age Group) | group | 1 | 40 | 0.009 | 0.9241464 |
| | ROI | 2 | 200 | 0.264 | 0.7679433 |
| | hemi | 1 | 200 | 10756.457 | 7.777e-176 |
| | group:ROI | 2 | 200 | 0.374 | 0.6885144 |
| | group:hemi | 1 | 200 | 0.028 | 0.8674489 |
| | ROI:hemi | 2 | 200 | 1.202 | 0.3027704 |
| | group:ROI:hemi | 2 | 200 | 1.066 | 0.3463284 |
| LMM: pRF centers Y-position (Age Group) | group | 1 | 40 | 0.011 | 0.9168957 |
| | ROI | 2 | 200 | 0.358 | 0.6994219 |
| | hemi | 1 | 200 | 0.683 | 0.4095495 |
| | group:ROI | 2 | 200 | 2.459 | 0.08813249 |
| | group:hemi | 1 | 200 | 0.274 | 0.6013173 |
| | ROI:hemi | 2 | 200 | 1.214 | 0.2991804 |
| | group:ROI:hemi | 2 | 200 | 1.152 | 0.3181497 |
| LMM: pRF centers Eccentricity (Age Group) | group | 1 | 40 | 0.050 | 0.8234434 |
| | ROI | 2 | 200 | 0.721 | 0.4874026 |
| | hemi | 1 | 200 | 22.907 | 3.299e-06 |
| | group:ROI | 2 | 200 | 1.911 | 0.1506917 |
| | group:hemi | 1 | 200 | 0.014 | 0.9044544 |
| | ROI:hemi | 2 | 200 | 0.140 | 0.8690739 |
| | group:ROI:hemi | 2 | 200 | 0.748 | 0.4747786 |
| LMM: pRF size (Age Group) | group | 1 | 40 | 0.154 | 0.6970016 |
| | ROI | 2 | 200 | 434.138 | 1.717e-73 |
| | hemi | 1 | 200 | 10.789 | 0.001205672 |
| | group:ROI | 2 | 200 | 0.334 | 0.7166424 |
| | group:hemi | 1 | 200 | 0.038 | 0.8461736 |
| | ROI:hemi | 2 | 200 | 8.805 | 0.0002164277 |
| | group:ROI:hemi | 2 | 200 | 1.016 | 0.3637664 |

**Supplementary Table 2: Relationship between pRF size v. eccentricity.**
Results of LMM (*pRF Size ~ Eccentricity x Age Group x ROI (V1/V2/V3) x Hemisphere + (1| participant)*) examining differences in pRF size across factors: eccentricity, age group, ROI, and hemisphere in early visual cortex (V1, V2, V3).

| Analysis | model term | df1 | df2 | F.ratio | p.value |
|---|---|---|---|---|---|
| LMM: Size v. Ecc Slopes | group | 1 | 33 | 0.018 | 0.8933011 |
| | ROI | 2 | 165 | 79.216 | 7.681e-25 |
| | hemi | 1 | 165 | 0.083 | 0.7733338 |
| | group:ROI | 2 | 165 | 0.694 | 0.5009582 |
| | group:hemi | 1 | 165 | 0.145 | 0.7034837 |
| | ROI:hemi | 2 | 165 | 2.689 | 0.07090125 |
| | group:ROI:hemi | 2 | 165 | 1.701 | 0.1856676 |
| LMM: Size v. Ecc Intercepts | group | 1 | 33 | 0.291 | 0.5930899 |
| | ROI | 2 | 165 | 158.376 | 4.069e-39 |
| | hemi | 1 | 165 | 11.113 | 0.001058557 |
| | group:ROI | 2 | 165 | 0.983 | 0.3764351 |
| | group:hemi | 1 | 165 | 0.055 | 0.8150063 |
| | ROI:hemi | 2 | 165 | 2.651 | 0.07358527 |
| | group:ROI:hemi | 2 | 165 | 0.267 | 0.7663083 |

**Supplementary Table 3: Participant counts for ROIs across hemispheres and age groups.** Number of adolescents (out of 15) and adults (out of 27) with left and right hemisphere ventral (top), lateral (middle), and dorsal (bottom) ROIs that had ten or more voxels modulated by the Toonotopy experiment. Rows with asterisks (*) denote regions derived from maximal probability maps.

| | Left Hemisphere | | Right Hemisphere | |
|---|---|---|---|---|
| ROI | adolescents | Adults | adolescents | Adults |
| IOG-faces | 6 | 16 | 11 | 20 |
| pFus-faces | 15 | 22 | 12 | 24 |
| mFus-faces | 14 | 20 | 12 | 21 |
| pOTS-words | 14 | 26 | 6 | 15 |
| mOTS-words | 8 | 12 | | |
| OTS-bodies | 11 | 14 | 11 | 19 |
| CoS-places | 14 | 24 | 14 | 25 |
| LOS-bodies | 14 | 25 | 15 | 24 |
| ITG-bodies | 11 | 21 | 13 | 23 |
| MTG-bodies* | 14 | 25 | 14 | 27 |
| pSTS-faces* | 10 | 11 | 4 | 9 |
| MOG-places | 15 | 23 | 15 | 25 |
| IPS-places | 15 | 25 | 15 | 25 |

**Supplementary Table 4: Differences in proportion of voxels with greater than 20% variance explained by the Toonotopy experiment.**

*Top:* Results of LMM (*proportion voxels with VE > 20% ~ Stream x Category x Age Group x Hemisphere +(1|participant)*) examining differences in proportion of voxels with >20% variance explained across factors: age group, stream, category, and hemisphere in ventral (excluding word-selective ROIs) and dorsal-lateral ROIs. *Bottom:* Results of ventral LMM (*proportion voxels with VE > 20% ~ Stream x Category x Age Group x Hemisphere +(1|participant)*) examining differences in proportion of voxels with >20% variance explained across factors: age group, category, and hemisphere in ventral ROIs.

| Analysis | model term | df1 | df2 | F.ratio | p.value |
|---|---|---|---|---|---|
| LMM: Proportion Voxels with Variance Explained > 20% | group | 1 | 55.19 | 3.076 | 0.08498666 |
| | stream | 1 | 689.98 | 13.222 | 0.0002973375 |
| | category | 2 | 683.89 | 31.444 | 8.625e-14 |
| | hemi | 1 | 689.08 | 14.143 | 0.0001838211 |
| | group:stream | 1 | 689.98 | 0.428 | 0.5133458 |
| | group:category | 2 | 683.89 | 4.635 | 0.01001403 |
| | group:hemi | 1 | 689.08 | 0.009 | 0.923237 |
| | stream:category | 2 | 683.20 | 72.143 | 3.755e-29 |
| | stream:hemi | 1 | 688.61 | 0.661 | 0.4164424 |
| | category:hemi | 2 | 682.80 | 0.302 | 0.7392237 |
| | group:stream:category | 2 | 683.20 | 2.491 | 0.08355253 |
| | group:stream:hemi | 1 | 688.61 | 0.335 | 0.5632079 |
| | group:category:hemi | 2 | 682.80 | 0.190 | 0.8269975 |
| | stream:category:hemi | 2 | 681.59 | 4.829 | 0.008272539 |
| | group:stream:category:hemi | 2 | 681.59 | 1.941 | 0.1443088 |
| Ventral LMM: Proportion Voxels with Variance Explained > 20% | group | 1 | 46.94 | 2.751 | 0.1038762 |
| | category | 3 | 355.73 | 36.727 | 1.063e-20 |
| | hemi | 1 | 359.21 | 9.374 | 0.002366371 |
| | group:category | 3 | 355.73 | 0.853 | 0.4655169 |
| | group:hemi | 1 | 359.21 | 1.069 | 0.3019352 |
| | category:hemi | 3 | 354.79 | 3.751 | 0.0112246 |
| | group:category:hemi | 3 | 354.79 | 0.879 | 0.4523456 |

**Supplementary Table 5: Differences in pRF x-position.**

*Analysis 1:* Results of LMM (*pRF x-position ~ Stream x Category x Age Group x Hemisphere +(1|participant)*) examining differences in pRF x-position explained across factors: age group, stream, category, and hemisphere in ventral (excluding word-selective ROIs) and dorsal-lateral ROIs. *Analysis 2:* Results of ventral LMM (*pRF x-position ~ Stream x Category x Age Group x Hemisphere +(1|participant)*) examining differences in pRF x-position explained across factors: age group, category, and hemisphere in ventral ROIs. *Analysis 3:* Same as Analysis 1 but with age as a continuous variable. *Analysis 4:* Same as Analysis 2 but with age as a continuous variable.

| Analysis | model term | df1 | df2 | F.ratio | p.value |
|---|---|---|---|---|---|
| LMM: pRF centers X-position (Age Group) | group | 1 | 55.09 | 0.571 | 0.4529406 |
| | stream | 1 | 690.13 | 2.411 | 0.1209122 |
| | category | 2 | 683.85 | 6.027 | 0.002542492 |
| | hemi | 1 | 688.85 | 2476.210 | 2.737e-230 |
| | group:stream | 1 | 690.13 | 2.184 | 0.1399435 |
| | group:category | 2 | 683.85 | 0.339 | 0.7124669 |
| | group:hemi | 1 | 688.85 | 5.177 | 0.0231882 |
| | stream:category | 2 | 682.95 | 8.247 | 0.000289024 |
| | stream:hemi | 1 | 688.54 | 43.983 | 6.7e-11 |
| | category:hemi | 2 | 682.49 | 13.702 | 1.465e-06 |
| | group:stream:category | 2 | 682.95 | 0.786 | 0.4562774 |
| | group:stream:hemi | 1 | 688.54 | 1.263 | 0.2615007 |
| | group:category:hemi | 2 | 682.49 | 0.098 | 0.906937 |
| | stream:category:hemi | 2 | 681.73 | 84.935 | 1.16e-33 |
| | group:stream:category:hemi | 2 | 681.73 | 1.385 | 0.2509704 |
| Ventral LMM: pRF centers X-position (Age Group) | group | 1 | 48.03 | 0.109 | 0.7431324 |
| | category | 3 | 355.15 | 1.410 | 0.2396151 |
| | hemi | 1 | 359.18 | 1730.317 | 2.121e-139 |
| | group:category | 3 | 355.15 | 0.317 | 0.8127501 |
| | group:hemi | 1 | 359.18 | 0.027 | 0.868421 |
| | category:hemi | 3 | 354.10 | 96.752 | 9.395e-46 |
| | group:category:hemi | 3 | 354.10 | 1.936 | 0.1233754 |
| LMM: pRF centers X-position (Age Continuous) | age | 1 | 56.42 | 0.201 | 0.6552662 |
| | stream | 1 | 691.49 | 4.119 | 0.04277641 |
| | category | 2 | 684.27 | 7.377 | 0.0006763734 |
| | hemi | 1 | 690.03 | 2554.665 | 3.792e-234 |
| | age:stream | 1 | 691.80 | 0.866 | 0.3524216 |
| | age:category | 2 | 684.15 | 0.197 | 0.8208409 |
| | age:hemi | 1 | 687.67 | 3.559 | 0.05964367 |
| | stream:category | 2 | 683.71 | 9.363 | 9.74e-05 |
| | stream:hemi | 1 | 688.14 | 43.114 | 1.018e-10 |
| | category:hemi | 2 | 682.96 | 13.954 | 1.149e-06 |
| | age:stream:category | 2 | 682.97 | 1.116 | 0.328225 |
| | age:stream:hemi | 1 | 690.17 | 0.002 | 0.9675697 |
| | age:category:hemi | 2 | 682.51 | 0.241 | 0.7855457 |
| | stream:category:hemi | 2 | 682.09 | 87.444 | 1.559e-34 |
| | age:stream:category:hemi | 2 | 682.12 | 0.311 | 0.7330919 |
| Ventral LMM: pRF centers X-position (Age Continuous) | age | 1 | 50.81 | 0.164 | 0.6874511 |
| | category | 3 | 355.80 | 1.467 | 0.2231668 |
| | hemi | 1 | 357.43 | 1876.079 | 2.764e-144 |
| | age:category | 3 | 355.49 | 0.368 | 0.7758979 |
| | age:hemi | 1 | 362.09 | 0.733 | 0.3926068 |
| | category:hemi | 3 | 354.45 | 98.269 | 2.641e-46 |
| | age:category:hemi | 3 | 354.53 | 1.742 | 0.1580918 |

**Supplementary Table 6: Differences in pRF y-position.**

*Analysis 1:* Results of LMM (*pRF y-position ~ Stream x Category x Age Group x Hemisphere +(1|participant)*) examining differences in pRF y-position explained across factors: age group, stream, category, and hemisphere in ventral (excluding word-selective ROIs) and dorsal-lateral ROIs. *Analysis 2:* Results of ventral LMM (*pRF y-position ~ Stream x Category x Age Group x Hemisphere +(1|participant)*) examining differences in pRF y-position explained across factors: age group, category, and hemisphere in ventral ROIs. *Analysis 3:* Same as Analysis 1 but with age as a continuous variable. *Analysis 4:* Same as Analysis 2 but with age as a continuous variable.

| Analysis | model term | df1 | df2 | F.ratio | p.value |
|---|---|---|---|---|---|
| LMM: pRF centers Y-position (Age Group) | group | 1 | 51.33 | 1.097 | 0.299898 |
| | stream | 1 | 688.08 | 3.331 | 0.06843386 |
| | category | 2 | 682.79 | 54.004 | 1.685e-22 |
| | hemi | 1 | 686.92 | 28.466 | 1.296e-07 |
| | group:stream | 1 | 688.08 | 1.560 | 0.2120987 |
| | group:category | 2 | 682.79 | 1.129 | 0.323957 |
| | group:hemi | 1 | 686.92 | 0.033 | 0.8553665 |
| | stream:category | 2 | 682.10 | 12.146 | 6.56e-06 |
| | stream:hemi | 1 | 686.63 | 0.431 | 0.5116942 |
| | category:hemi | 2 | 681.69 | 3.437 | 0.03271604 |
| | group:stream:category | 2 | 682.10 | 0.632 | 0.5317993 |
| | group:stream:hemi | 1 | 686.63 | 0.154 | 0.6951485 |
| | group:category:hemi | 2 | 681.69 | 0.117 | 0.8898775 |
| | stream:category:hemi | 2 | 681.09 | 2.158 | 0.1163485 |
| | group:stream:category:hemi | 2 | 681.09 | 0.416 | 0.6595351 |
| Ventral LMM: pRF centers Y-position (Age Group) | group | 1 | 46.43 | 0.277 | 0.6010871 |
| | category | 3 | 354.27 | 54.076 | 8.423e-29 |
| | hemi | 1 | 357.77 | 50.808 | 5.661e-12 |
| | group:category | 3 | 354.27 | 0.155 | 0.9261724 |
| | group:hemi | 1 | 357.77 | 0.059 | 0.8078982 |
| | category:hemi | 3 | 353.41 | 3.330 | 0.01974629 |
| | group:category:hemi | 3 | 353.41 | 0.262 | 0.8528185 |
| LMM: pRF centers Y-position (Age Continuous) | age | 1 | 52.56 | 0.947 | 0.3348539 |
| | stream | 1 | 689.44 | 2.369 | 0.1242039 |
| | category | 2 | 683.20 | 58.242 | 4.413e-24 |
| | hemi | 1 | 688.06 | 29.873 | 6.457e-08 |
| | age:stream | 1 | 689.55 | 3.406 | 0.06537624 |
| | age:category | 2 | 683.08 | 0.742 | 0.47649 |
| | age:hemi | 1 | 686.05 | 0.050 | 0.8227482 |
| | stream:category | 2 | 682.77 | 13.982 | 1.119e-06 |
| | stream:hemi | 1 | 686.52 | 0.324 | 0.5694836 |
| | category:hemi | 2 | 682.09 | 3.289 | 0.03787949 |
| | age:stream:category | 2 | 682.16 | 0.783 | 0.4575578 |
| | age:stream:hemi | 1 | 688.01 | 0.157 | 0.6916687 |
| | age:category:hemi | 2 | 681.74 | 0.325 | 0.722527 |
| | stream:category:hemi | 2 | 681.40 | 1.810 | 0.1645023 |
| | age:stream:category:hemi | 2 | 681.44 | 1.026 | 0.3591414 |
| Ventral LMM: pRF centers Y-position (Age Continuous) | age | 1 | 48.74 | 0.007 | 0.9350167 |
| | category | 3 | 354.77 | 56.969 | 4.407e-30 |
| | hemi | 1 | 356.16 | 53.813 | 1.502e-12 |
| | age:category | 3 | 354.53 | 0.064 | 0.9790183 |
| | age:hemi | 1 | 360.19 | 0.079 | 0.7792094 |
| | category:hemi | 3 | 353.63 | 3.784 | 0.01074705 |
| | age:category:hemi | 3 | 353.76 | 0.505 | 0.6793987 |

**Supplementary Table 7: Differences in pRF eccentricity.**

*Analysis 1:* Results of LMM (*Eccentricity ~ Stream x Category x Age Group x Hemisphere +(1|participant)*) examining differences in pRF eccentricity explained across factors: age group, stream, category, and hemisphere in ventral (excluding word-selective ROIs) and dorsal-lateral ROIs. *Analysis 2:* Results of ventral LMM (*Eccentricity ~ Stream x Category x Age Group x Hemisphere +(1|participant)*) examining differences in pRF eccentricity explained across factors: age group, category, and hemisphere in ventral ROIs. *Analysis 3:* Same as Analysis 1 but with age as a continuous variable. *Analysis 4:* Same as Analysis 2 but with age as a continuous variable.

| Analysis | model term | df1 | df2 | F.ratio | p.value |
|---|---|---|---|---|---|
| LMM: pRF centers Eccentricity (Age Group) | group | 1 | 61.32 | 0.387 | 0.5364031 |
| | stream | 1 | 693.18 | 84.616 | 4.185e-19 |
| | category | 2 | 685.49 | 6.213 | 0.002117335 |
| | hemi | 1 | 691.78 | 58.604 | 6.501e-14 |
| | group:stream | 1 | 693.18 | 0.107 | 0.7441892 |
| | group:category | 2 | 685.49 | 1.269 | 0.2817833 |
| | group:hemi | 1 | 691.78 | 0.089 | 0.7657363 |
| | stream:category | 2 | 684.25 | 106.505 | 5.379e-41 |
| | stream:hemi | 1 | 691.46 | 1.426 | 0.2327725 |
| | category:hemi | 2 | 683.74 | 7.575 | 0.0005572182 |
| | group:stream:category | 2 | 684.25 | 0.330 | 0.7191531 |
| | group:stream:hemi | 1 | 691.46 | 0.477 | 0.4902092 |
| | group:category:hemi | 2 | 683.74 | 0.094 | 0.9103782 |
| | stream:category:hemi | 2 | 682.72 | 10.701 | 2.654e-05 |
| | group:stream:category:hemi | 2 | 682.72 | 0.228 | 0.7964207 |
| Ventral LMM: pRF centers Eccentricity (Age Group) | group | 1 | 54.52 | 0.372 | 0.5446127 |
| | category | 3 | 358.50 | 84.591 | 2.103e-41 |
| | hemi | 1 | 364.21 | 21.269 | 5.534e-06 |
| | group:category | 3 | 358.50 | 1.185 | 0.3151562 |
| | group:hemi | 1 | 364.21 | 0.004 | 0.9523363 |
| | category:hemi | 3 | 356.71 | 5.069 | 0.001893166 |
| | group:category:hemi | 3 | 356.71 | 0.447 | 0.7195181 |
| LMM: pRF centers Eccentricity (Age Continuous) | age | 1 | 63.42 | 0.716 | 0.4007952 |
| | stream | 1 | 694.77 | 90.471 | 3.004e-20 |
| | category | 2 | 686.09 | 5.280 | 0.005300809 |
| | hemi | 1 | 693.26 | 64.064 | 5.076e-15 |
| | age:stream | 1 | 695.44 | 0.747 | 0.3878589 |
| | age:category | 2 | 685.99 | 1.891 | 0.1516833 |
| | age:hemi | 1 | 690.39 | 0.094 | 0.7591892 |
| | stream:category | 2 | 685.29 | 109.945 | 3.87e-42 |
| | stream:hemi | 1 | 690.82 | 2.156 | 0.1424806 |
| | category:hemi | 2 | 684.45 | 8.341 | 0.0002635317 |
| | age:stream:category | 2 | 684.36 | 0.187 | 0.8297307 |
| | age:stream:hemi | 1 | 693.81 | 0.512 | 0.4745478 |
| | age:category:hemi | 2 | 683.81 | 0.006 | 0.9936383 |
| | stream:category:hemi | 2 | 683.30 | 10.815 | 2.376e-05 |
| | age:stream:category:hemi | 2 | 683.30 | 0.103 | 0.9023179 |
| Ventral LMM: pRF centers Eccentricity (Age Continuous) | age | 1 | 58.35 | 0.756 | 0.3882737 |
| | category | 3 | 359.61 | 85.788 | 6.913e-42 |
| | hemi | 1 | 361.99 | 23.888 | 1.537e-06 |
| | age:category | 3 | 359.00 | 1.362 | 0.254283 |
| | age:hemi | 1 | 368.36 | 0.396 | 0.5296257 |
| | category:hemi | 3 | 357.56 | 4.873 | 0.002467412 |
| | age:category:hemi | 3 | 357.26 | 0.106 | 0.9562657 |

**Supplementary Table 8: Differences in pRF size.**

*Analysis 1:* Results of LMM (*pRF size ~ Stream x Category x Age Group x Hemisphere +(1|participant)*) examining differences in pRF size explained across factors: age group, stream, category, and hemisphere in ventral (excluding word-selective ROIs) and dorsal-lateral ROIs. *Analysis 2:* Results of ventral LMM (*pRF size ~ Stream x Category x Age Group x Hemisphere +(1|participant)*) examining differences in pRF size explained across factors: age group, category, and hemisphere in ventral ROIs. *Analysis 3:* Same as Analysis 1 but with age as a continuous variable. *Analysis 4:* Same as Analysis 2 but with age as a continuous variable.

| Analysis | model term | df1 | df2 | F.ratio | p.value |
|---|---|---|---|---|---|
| LMM: pRF Size (Age Group) | group | 1 | 64.20 | 5.621 | 0.0207624 |
| | stream | 1 | 694.48 | 35.637 | 3.791e-09 |
| | category | 2 | 686.21 | 21.692 | 7.333e-10 |
| | hemi | 1 | 693.04 | 1.707 | 0.1918125 |
| | group:stream | 1 | 694.48 | 3.287 | 0.07024347 |
| | group:category | 2 | 686.21 | 1.229 | 0.2933021 |
| | group:hemi | 1 | 693.04 | 9.828 | 0.001791719 |
| | stream:category | 2 | 684.82 | 25.675 | 1.769e-11 |
| | stream:hemi | 1 | 692.73 | 0.862 | 0.3536387 |
| | category:hemi | 2 | 684.29 | 0.529 | 0.5893665 |
| | group:stream:category | 2 | 684.82 | 1.478 | 0.2287811 |
| | group:stream:hemi | 1 | 692.73 | 4.060 | 0.04429823 |
| | group:category:hemi | 2 | 684.29 | 2.948 | 0.05310739 |
| | stream:category:hemi | 2 | 683.16 | 0.574 | 0.5633903 |
| | group:stream:category:hemi | 2 | 683.16 | 3.615 | 0.02743174 |
| Ventral LMM: pRF Size (Age Group) | group | 1 | 55.86 | 3.943 | 0.05197398 |
| | category | 3 | 359.16 | 19.069 | 1.699e-11 |
| | hemi | 1 | 365.14 | 1.313 | 0.2525911 |
| | group:category | 3 | 359.16 | 0.635 | 0.5927997 |
| | group:hemi | 1 | 365.14 | 2.502 | 0.1145317 |
| | category:hemi | 3 | 357.22 | 0.341 | 0.7954864 |
| | group:category:hemi | 3 | 357.22 | 1.573 | 0.1955183 |
| LMM: pRF Size (Age Continuous) | age | 1 | 65.58 | 4.232 | 0.04365917 |
| | stream | 1 | 695.71 | 33.297 | 1.191e-08 |
| | category | 2 | 686.63 | 20.448 | 2.366e-09 |
| | hemi | 1 | 694.20 | 0.510 | 0.4754093 |
| | age:stream | 1 | 696.47 | 3.785 | 0.05212508 |
| | age:category | 2 | 686.54 | 1.062 | 0.3464486 |
| | age:hemi | 1 | 691.19 | 7.370 | 0.006797113 |
| | stream:category | 2 | 685.75 | 24.906 | 3.617e-11 |
| | stream:hemi | 1 | 691.60 | 0.366 | 0.5453344 |
| | category:hemi | 2 | 684.90 | 0.403 | 0.6686371 |
| | age:stream:category | 2 | 684.77 | 1.734 | 0.1774192 |
| | age:stream:hemi | 1 | 694.88 | 3.926 | 0.04793761 |
| | age:category:hemi | 2 | 684.19 | 2.538 | 0.07976646 |
| | stream:category:hemi | 2 | 683.66 | 0.208 | 0.8125326 |
| | age:stream:category:hemi | 2 | 683.65 | 3.312 | 0.03703415 |
| Ventral LMM: pRF Size (Age Continuous) | age | 1 | 59.35 | 1.468 | 0.2305008 |
| | category | 3 | 360.12 | 18.135 | 5.565e-11 |
| | hemi | 1 | 362.60 | 0.506 | 0.4773561 |
| | age:category | 3 | 359.46 | 0.146 | 0.9323338 |
| | age:hemi | 1 | 369.12 | 0.816 | 0.3670238 |
| | category:hemi | 3 | 357.98 | 0.460 | 0.7101169 |
| | age:category:hemi | 3 | 357.62 | 1.192 | 0.312757 |

**Supplementary Table 9: Differences in VFC FWHM.**

*Analysis 1:* Results of LMM (*FWHM ~ Stream x Category x Age Group x Hemisphere +(1|participant)*) examining differences in VFC FWHM explained across factors: age group, stream, category, and hemisphere in ventral (excluding word-selective ROIs) and dorsal-lateral ROIs. *Analysis 2:* Results of ventral LMM (*FWHM ~ Stream x Category x Age Group x Hemisphere +(1|participant)*) examining differences in VFC FWHM explained across factors: age group, category, and hemisphere in ventral ROIs. *Analysis 3:* Same as Analysis 1 but with age as a continuous variable. *Analysis 4:* Same as Analysis 2 but with age as a continuous variable.

| Analysis | model term | df1 | df2 | F.ratio | p.value |
|---|---|---|---|---|---|
| LMM: Total FWHM (Age Group) | group | 1 | 65.46 | 1.837 | 0.1799715 |
| | stream | 1 | 694.72 | 7.024 | 0.00822665 |
| | category | 2 | 686.50 | 4.094 | 0.01707638 |
| | hemi | 1 | 693.82 | 3.636 | 0.05695543 |
| | group:stream | 1 | 694.72 | 4.782 | 0.02909658 |
| | group:category | 2 | 686.50 | 0.883 | 0.4139856 |
| | group:hemi | 1 | 693.82 | 3.248 | 0.0719624 |
| | stream:category | 2 | 685.38 | 14.525 | 6.638e-07 |
| | stream:hemi | 1 | 693.29 | 0.227 | 0.6337889 |
| | category:hemi | 2 | 684.95 | 2.181 | 0.113698 |
| | group:stream:category | 2 | 685.38 | 5.996 | 0.002620842 |
| | group:stream:hemi | 1 | 693.29 | 7.148 | 0.007680015 |
| | group:category:hemi | 2 | 684.95 | 0.859 | 0.4242138 |
| | stream:category:hemi | 2 | 683.10 | 0.624 | 0.5359071 |
| | group:stream:category:hemi | 2 | 683.10 | 3.513 | 0.03034913 |
| Ventral LMM: Total FWHM (Age Group) | group | 1 | 52.03 | 0.013 | 0.9103476 |
| | category | 3 | 358.41 | 13.639 | 1.914e-08 |
| | hemi | 1 | 363.33 | 0.781 | 0.3772795 |
| | group:category | 3 | 358.41 | 1.536 | 0.2049264 |
| | group:hemi | 1 | 363.33 | 0.338 | 0.5614568 |
| | category:hemi | 3 | 356.91 | 1.327 | 0.2654512 |
| | group:category:hemi | 3 | 356.91 | 0.767 | 0.5131287 |
| LMM: Total FWHM (Age Continuous) | group | 1 | 65.46 | 1.837 | 0.1799715 |
| | stream | 1 | 694.72 | 7.024 | 0.00822665 |
| | category | 2 | 686.50 | 4.094 | 0.01707638 |
| | hemi | 1 | 693.82 | 3.636 | 0.05695543 |
| | group:stream | 1 | 694.72 | 4.782 | 0.02909658 |
| | group:category | 2 | 686.50 | 0.883 | 0.4139856 |
| | group:hemi | 1 | 693.82 | 3.248 | 0.0719624 |
| | stream:category | 2 | 685.38 | 14.525 | 6.638e-07 |
| | stream:hemi | 1 | 693.29 | 0.227 | 0.6337889 |
| | category:hemi | 2 | 684.95 | 2.181 | 0.113698 |
| | group:stream:category | 2 | 685.38 | 5.996 | 0.002620842 |
| | group:stream:hemi | 1 | 693.29 | 7.148 | 0.007680015 |
| | group:category:hemi | 2 | 684.95 | 0.859 | 0.4242138 |
| | stream:category:hemi | 2 | 683.10 | 0.624 | 0.5359071 |
| | group:stream:category:hemi | 2 | 683.10 | 3.513 | 0.03034913 |
| Ventral LMM: Total FWHM (Age Continuous) | group | 1 | 52.03 | 0.013 | 0.9103476 |
| | category | 3 | 358.41 | 13.639 | 1.914e-08 |
| | hemi | 1 | 363.33 | 0.781 | 0.3772795 |
| | group:category | 3 | 358.41 | 1.536 | 0.2049264 |
| | group:hemi | 1 | 363.33 | 0.338 | 0.5614568 |
| | category:hemi | 3 | 356.91 | 1.327 | 0.2654512 |
| | group:category:hemi | 3 | 356.91 | 0.767 | 0.5131287 |

**Supplementary Table 10. Visual field coverage full-width half max percentages.**

Mean percentage of FWHM (± standard deviation) in the upper contralateral (UC), lower contralateral (LC), upper ipsilateral (UI), lower ipsilateral (LI), and central 5° in the left (lh) and right (rh) hemisphere in adolescents and adults.

| | | UC | | LC | | LI | | UI | | <5° | |
|---|---|---|---|---|---|---|---|---|---|---|---|
| **ROI** | **Group** | **lh** | **rh** | **lh** | **rh** | **lh** | **rh** | **lh** | **rh** | **lh** | **rh** |
| IOG-faces | adolescents | 40 ± 36 | 27 ± 32 | 54 ± 39 | 58 ± 29 | 2 ± 2 | 13 ± 18 | 4 ± 9 | 2 ± 4 | 33 ± 41 | 54 ± 30 |
| IOG-faces | Adults | 49 ± 31 | 37 ± 26 | 48 ± 32 | 50 ± 26 | 1 ± 2 | 7 ± 11 | 2 ± 5 | 5 ± 9 | 27 ± 37 | 39 ± 27 |
| pFus-faces | adolescents | 44 ± 29 | 30 ± 27 | 49 ± 25 | 54 ± 27 | 6 ± 15 | 10 ± 11 | 1 ± 2 | 5 ± 7 | 28 ± 21 | 42 ± 23 |
| pFus-faces | Adults | 47 ± 29 | 35 ± 24 | 43 ± 27 | 46 ± 24 | 6 ± 18 | 12 ± 14 | 3 ± 10 | 8 ± 9 | 27 ± 24 | 40 ± 19 |
| mFus-faces | adolescents | 50 ± 13 | 34 ± 23 | 43 ± 15 | 36 ± 27 | 3 ± 4 | 18 ± 27 | 4 ± 7 | 12 ± 11 | 39 ± 22 | 58 ± 27 |
| mFus-faces | Adults | 40 ± 18 | 29 ± 19 | 48 ± 21 | 50 ± 22 | 6 ± 10 | 14 ± 13 | 6 ± 9 | 8 ± 7 | 34 ± 20 | 51 ± 20 |
| pOTS-words | adolescents | 34 ± 29 | 17 ± 26 | 56 ± 29 | 74 ± 23 | 7 ± 14 | 9 ± 13 | 3 ± 7 | 0 ± 0 | 36 ± 21 | 24 ± 16 |
| pOTS-words | Adults | 42 ± 29 | 22 ± 22 | 52 ± 29 | 71 ± 25 | 3 ± 7 | 5 ± 12 | 2 ± 5 | 2 ± 5 | 29 ± 23 | 33 ± 26 |
| mOTS-words | adolescents | 26 ± 25 | | 59 ± 23 | | 12 ± 21 | | 2 ± 3 | | 22 ± 29 | |
| mOTS-words | Adults | 44 ± 26 | | 52 ± 28 | | 1 ± 2 | | 3 ± 8 | | 34 ± 30 | |
| OTS-bodies | adolescents | 34 ± 23 | 24 ± 26 | 59 ± 28 | 57 ± 21 | 3 ± 4 | 17 ± 16 | 4 ± 10 | 3 ± 4 | 32 ± 26 | 38 ± 25 |
| OTS-bodies | Adults | 22 ± 25 | 17 ± 23 | 67 ± 30 | 56 ± 29 | 7 ± 10 | 24 ± 29 | 4 ± 10 | 2 ± 5 | 32 ± 28 | 30 ± 20 |
| CoS-places | adolescents | 72 ± 18 | 46 ± 15 | 27 ± 18 | 49 ± 15 | 0 ± 0 | 3 ± 4 | 0 ± 1 | 2 ± 5 | 6 ± 8 | 20 ± 11 |
| CoS-places | Adults | 72 ± 20 | 51 ± 26 | 27 ± 20 | 42 ± 23 | 0 ± 2 | 4 ± 8 | 1 ± 3 | 3 ± 6 | 8 ± 10 | 18 ± 11 |
| LOS-limbs | adolescents | 33 ± 21 | 32 ± 21 | 65 ± 23 | 65 ± 24 | 1 ± 3 | 2 ± 6 | 1 ± 3 | 2 ± 5 | 6 ± 11 | 10 ± 11 |
| LOS-limbs | Adults | 37 ± 23 | 29 ± 17 | 62 ± 25 | 65 ± 21 | 1 ± 2 | 3 ± 8 | 1 ± 4 | 2 ± 5 | 12 ± 18 | 16 ± 12 |
| ITG-limbs | adolescents | 36 ± 27 | 19 ± 22 | 55 ± 32 | 77 ± 23 | 4 ± 8 | 3 ± 6 | 5 ± 10 | 1 ± 2 | 20 ± 19 | 12 ± 12 |
| ITG-limbs | Adults | 39 ± 32 | 21 ± 24 | 56 ± 33 | 67 ± 27 | 2 ± 7 | 9 ± 13 | 2 ± 6 | 2 ± 5 | 9 ± 11 | 20 ± 20 |
| MTG-limbs | adolescents | 50 ± 31 | 29 ± 21 | 36 ± 30 | 48 ± 27 | 5 ± 9 | 12 ± 11 | 9 ± 15 | 10 ± 11 | 15 ± 15 | 21 ± 14 |
| MTG-limbs | Adults | 39 ± 27 | 25 ± 23 | 47 ± 31 | 54 ± 29 | 6 ± 8 | 14 ± 16 | 8 ± 10 | 8 ± 13 | 20 ± 13 | 20 ± 14 |
| pSTS-faces | adolescents | 40 ± 30 | 38 ± 23 | 55 ± 29 | 28 ± 7 | 4 ± 9 | 17 ± 17 | 2 ± 4 | 16 ± 9 | 9 ± 15 | 39 ± 13 |
| pSTS-faces | Adults | 55 ± 36 | 43 ± 37 | 37 ± 35 | 30 ± 34 | 4 ± 10 | 13 ± 24 | 4 ± 7 | 15 ± 23 | 9 ± 13 | 29 ± 37 |
| MOG-places | adolescents | 34 ± 29 | 27 ± 21 | 66 ± 29 | 73 ± 21 | 0 ± 0 | 0 ± 1 | 0 ± 0 | 0 ± 0 | 13 ± 12 | 15 ± 8 |
| MOG-places | Adults | 28 ± 27 | 23 ± 28 | 71 ± 27 | 73 ± 28 | 0 ± 1 | 4 ± 7 | 0 ± 0 | 1 ± 4 | 11 ± 11 | 18 ± 13 |
| IPS-places | adolescents | 67 ± 24 | 47 ± 18 | 32 ± 24 | 49 ± 17 | 0 ± 0 | 2 ± 7 | 1 ± 3 | 2 ± 4 | 11 ± 14 | 15 ± 14 |
| IPS-places | Adults | 69 ± 26 | 46 ± 35 | 28 ± 27 | 50 ± 33 | 0 ± 1 | 3 ± 8 | 2 ± 9 | 1 ± 3 | 9 ± 9 | 16 ± 13 |

**Supplementary Table 11: Differences in category selectivity - mean t-value.**
*Analysis 1:* Results of LMM (*Mean T-Value ~ Stream x Category x Age Group x Hemisphere +(1|participant)*) examining differences in category selectivity (mean t-value) explained across factors: age group, stream, category, and hemisphere in ventral (excluding word-selective ROIs) and dorsal-lateral ROIs. *Analysis 2:* Results of ventral LMM (*Mean T-Value ~ Stream x Category x Age Group x Hemisphere +(1|participant)*) examining differences in category selectivity (mean t-value) explained across factors: age group, category, and hemisphere in ventral ROIs. *Analysis 3:* Same as Analysis 1 but with age as a continuous variable. *Analysis 4:* Same as Analysis 2 but with age as a continuous variable.

| Analysis | model term | df1 | df2 | F.ratio | p.value |
|---|---|---|---|---|---|
| LMM: Category Selectivity - Mean T-Value (Age Group) | group | 1 | 41.39 | 3.649 | 0.06304484 |
| | stream | 1 | 748.43 | 8.321 | 0.00403196 |
| | category | 2 | 746.92 | 151.266 | 6.985e-56 |
| | hemi | 1 | 746.72 | 16.336 | 5.856e-05 |
| | group:stream | 1 | 748.43 | 2.911 | 0.08841133 |
| | group:category | 2 | 746.92 | 15.575 | 2.36e-07 |
| | group:hemi | 1 | 746.72 | 2.155 | 0.1424888 |
| | stream:category | 2 | 747.44 | 38.634 | 1.079e-16 |
| | stream:hemi | 1 | 746.95 | 0.272 | 0.6019996 |
| | category:hemi | 2 | 746.63 | 0.622 | 0.5368838 |
| | group:stream:category | 2 | 747.44 | 1.656 | 0.191631 |
| | group:stream:hemi | 1 | 746.95 | 0.517 | 0.4725215 |
| | group:category:hemi | 2 | 746.63 | 0.628 | 0.5338558 |
| | stream:category:hemi | 2 | 746.65 | 0.283 | 0.7533263 |
| | group:stream:category:hemi | 2 | 746.65 | 0.133 | 0.875892 |
| Ventral LMM: Category Selectivity - Mean T-Value (Age Group) | group | 1 | 44.48 | 3.380 | 0.07269208 |
| | category | 3 | 358.77 | 33.336 | 4.985e-19 |
| | hemi | 1 | 361.51 | 4.293 | 0.03897634 |
| | group:category | 3 | 358.77 | 5.098 | 0.001818502 |
| | group:hemi | 1 | 361.51 | 2.172 | 0.1413857 |
| | category:hemi | 3 | 357.48 | 1.799 | 0.1469729 |
| | group:category:hemi | 3 | 357.48 | 1.165 | 0.3228932 |
| LMM: Category Selectivity - Mean T-Value (Age Continuous) | age | 1 | 41.56 | 5.177 | 0.02811097 |
| | stream | 1 | 748.46 | 6.675 | 0.009965578 |
| | category | 2 | 747.01 | 139.270 | 3.931e-52 |
| | hemi | 1 | 746.60 | 14.457 | 0.0001551382 |
| | age:stream | 1 | 748.94 | 3.267 | 0.07110236 |
| | age:category | 2 | 747.14 | 19.314 | 6.632e-09 |
| | age:hemi | 1 | 746.94 | 3.042 | 0.08154305 |
| | stream:category | 2 | 747.33 | 38.909 | 8.419e-17 |
| | stream:hemi | 1 | 746.83 | 0.078 | 0.7795987 |
| | category:hemi | 2 | 746.75 | 1.050 | 0.3505715 |
| | age:stream:category | 2 | 747.38 | 2.944 | 0.05325067 |
| | age:stream:hemi | 1 | 747.16 | 1.374 | 0.2415592 |
| | age:category:hemi | 2 | 746.61 | 0.763 | 0.4665975 |
| | stream:category:hemi | 2 | 746.74 | 0.415 | 0.6602051 |
| | age:stream:category:hemi | 2 | 746.67 | 0.139 | 0.8702337 |
| Ventral LMM: Category Selectivity - Mean T-Value (Age Continuous) | age | 1 | 47.41 | 4.963 | 0.03068469 |
| | category | 3 | 358.98 | 29.109 | 7.206e-17 |
| | hemi | 1 | 360.56 | 3.150 | 0.07677821 |
| | age:category | 3 | 359.14 | 5.047 | 0.001948184 |
| | age:hemi | 1 | 363.33 | 1.201 | 0.2739205 |
| | category:hemi | 3 | 357.63 | 2.729 | 0.04384259 |
| | age:category:hemi | 3 | 357.50 | 0.855 | 0.4644582 |

**Supplementary Table 12: Development of category selective and ROI size.**

As a complementary analysis to the analyses in Supplementary Table 11, we evaluated the spatial extent of category selectivity. We measured the size of each ROI by quantifying the number of selective voxels (t > 3, for category of interest) in each region and performed an LMM to assess significant differences across age groups, streams, categories and hemispheres. *Analysis 1:* Results of LMM (*ROI Size ~ Stream x Category x Age Group x Hemisphere +(1|participant)*) examining differences in category ROI size explained across factors: age group, stream, category, and hemisphere in ventral (excluding word-selective ROIs) and dorsal-lateral ROIs. *Analysis 2:* Results of ventral LMM (*ROI Size ~ Stream x Category x Age Group x Hemisphere +(1|participant)*) examining differences in category ROI size explained across factors: age group, category, and hemisphere in ventral ROIs. *Analysis 3:* Same as Analysis 1 but with age as a continuous variable. *Analysis 4:* Same as Analysis 2 but with age as a continuous variable. Like with the mean t-values, we found a significant group by category effect, indicating that the size of ROIs differentially develops across visual categories.

| Analysis | model term | df1 | df2 | F.ratio | p.value |
|---|---|---|---|---|---|
| LMM: ROI Size (Age Group) | group | 1 | 46.77 | 0.012 | 0.9128761 |
| | stream | 1 | 781.27 | 9.632 | 0.001980754 |
| | category | 2 | 778.79 | 98.477 | 7.453e-39 |
| | hemi | 1 | 778.44 | 9.450 | 0.002184879 |
| | group:stream | 1 | 781.27 | 0.045 | 0.8324844 |
| | group:category | 2 | 778.79 | 5.905 | 0.002849194 |
| | group:hemi | 1 | 778.44 | 0.028 | 0.8664404 |
| | stream:category | 2 | 778.56 | 20.548 | 2.011e-09 |
| | stream:hemi | 1 | 779.03 | 1.083 | 0.2982953 |
| | category:hemi | 2 | 777.75 | 0.523 | 0.5928942 |
| | group:stream:category | 2 | 778.56 | 0.766 | 0.4654435 |
| | group:stream:hemi | 1 | 779.03 | 0.536 | 0.464113 |
| | group:category:hemi | 2 | 777.75 | 0.621 | 0.5376763 |
| | stream:category:hemi | 2 | 778.20 | 0.237 | 0.7887739 |
| | group:stream:category:hemi | 2 | 778.20 | 0.706 | 0.4937521 |
| Ventral LMM: ROI Size (Age Group) | group | 1 | 53.15 | 1.276 | 0.2637247 |
| | category | 3 | 386.02 | 34.519 | 8.748e-20 |
| | hemi | 1 | 389.55 | 1.084 | 0.298432 |
| | group:category | 3 | 386.02 | 4.283 | 0.005442802 |
| | group:hemi | 1 | 389.55 | 2.808 | 0.09456862 |
| | category:hemi | 3 | 384.71 | 5.967 | 0.0005525804 |
| | group:category:hemi | 3 | 384.71 | 2.220 | 0.08537581 |
| LMM: ROI Size (Age Continuous) | age | 1 | 47.77 | 0.007 | 0.9358654 |
| | stream | 1 | 782.14 | 9.844 | 0.001767452 |
| | category | 2 | 779.22 | 91.885 | 1.482e-36 |
| | hemi | 1 | 778.22 | 9.658 | 0.00195399 |
| | age:stream | 1 | 783.66 | 0.062 | 0.8029798 |
| | age:category | 2 | 778.86 | 3.162 | 0.04286842 |
| | age:hemi | 1 | 778.78 | 0.060 | 0.806741 |
| | stream:category | 2 | 778.83 | 23.753 | 9.696e-11 |
| | stream:hemi | 1 | 778.86 | 0.754 | 0.3854444 |
| | category:hemi | 2 | 778.20 | 0.269 | 0.7644172 |
| | age:stream:category | 2 | 778.66 | 0.349 | 0.7053723 |
| | age:stream:hemi | 1 | 779.81 | 0.366 | 0.5453664 |
| | age:category:hemi | 2 | 777.66 | 0.480 | 0.6192021 |
| | stream:category:hemi | 2 | 778.36 | 0.315 | 0.7296817 |
| | age:stream:category:hemi | 2 | 778.61 | 0.420 | 0.6569299 |
| Ventral LMM: ROI Size (Age Continuous) | age | 1 | 57.14 | 1.562 | 0.2164373 |
| | category | 3 | 386.85 | 31.288 | 3.96e-18 |
| | hemi | 1 | 388.43 | 0.422 | 0.5163726 |
| | age:category | 3 | 386.30 | 1.876 | 0.1330115 |
| | age:hemi | 1 | 392.58 | 2.067 | 0.1512864 |
| | category:hemi | 3 | 385.40 | 7.355 | 8.35e-05 |
| | age:category:hemi | 3 | 385.12 | 1.748 | 0.1567066 |

**Supplementary Table 13: Relationship between FWHM v. category selectivity (mean t-value).**

*Analysis 1:* Results of LMM (*Mean T-Value ~ FWHM x Stream x Category x Hemisphere +(1|participant)*) examining the relationship between category selectivity and FWHM (factors: FWHM, stream, category, and hemisphere) in ventral (excluding word-selective ROIs) and dorsal-lateral ROIs. *Analysis 2:* Results of ventral LMM (*Mean T-Value ~ FWHM x Category x Hemisphere +(1|participant)*) examining differences in between category selectivity and FWHM (factors: FWHM, category, and hemisphere) in ventral ROIs.

| Analysis | model term | df1 | df2 | F.ratio | p.value |
|---|---|---|---|---|---|
| LMM: Category Selectivity v FWHM (No Age) | fwhm | 1 | 628.47 | 4.665 | 0.03116602 |
| | stream | 1 | 621.60 | 3.541 | 0.06032356 |
| | category | 2 | 620.53 | 69.901 | 4.189e-28 |
| | hemi | 1 | 621.19 | 14.137 | 0.0001860192 |
| | fwhm:stream | 1 | 625.18 | 1.123 | 0.2896035 |
| | fwhm:category | 2 | 623.48 | 1.272 | 0.2810506 |
| | fwhm:hemi | 1 | 626.10 | 0.253 | 0.6151244 |
| | stream:category | 2 | 620.76 | 25.822 | 1.69e-11 |
| | stream:hemi | 1 | 620.37 | 0.398 | 0.5282362 |
| | category:hemi | 2 | 619.81 | 2.155 | 0.1167477 |
| | fwhm:stream:category | 2 | 624.30 | 5.030 | 0.006807533 |
| | fwhm:stream:hemi | 1 | 625.12 | 0.264 | 0.6078787 |
| | fwhm:category:hemi | 2 | 622.42 | 0.621 | 0.5379904 |
| | stream:category:hemi | 2 | 619.32 | 0.251 | 0.7783051 |
| | fwhm:stream:category:hemi | 2 | 623.68 | 0.929 | 0.3953425 |
| Ventral LMM: Category Selectivity v FWHM (No Age) | fwhm | 1 | 323.53 | 15.633 | 9.451e-05 |
| | category | 3 | 311.17 | 18.783 | 3.19e-11 |
| | hemi | 1 | 314.76 | 4.098 | 0.04377023 |
| | fwhm:category | 3 | 316.64 | 2.540 | 0.05650642 |
| | fwhm:hemi | 1 | 322.29 | 0.712 | 0.3994799 |
| | category:hemi | 3 | 312.87 | 2.595 | 0.05260194 |
| | fwhm:category:hemi | 3 | 319.37 | 0.962 | 0.4110887 |
